# 5’UTR-mediated retention of eIF3 on 80S ribosomes promotes co-translational folding of ER membrane proteins

**DOI:** 10.1101/2025.01.19.633751

**Authors:** Baochun Han, Siqiong Zhang, Haoran Duan, Yurui Zhang, Binbin Yu, Weimin Lin, Yabin Cheng, Dieter A. Wolf

## Abstract

Co-translational folding of nascent polypeptides is essential for protein function and cellular homeostasis. Ribosome-associated chaperones assist in this process, but coordination between their recruitment and translation initiation remains poorly understood. We report here that specific binding sites for eukaryotic translation initiation factor eIF3 within the 5’ untranslated regions (5’UTRs) of mRNAs promote its retention on 80S ribosomes during the synthesis of select endoplasmic reticulum (ER) membrane proteins. Disruption of these eIF3 binding sites leads to misfolding and sequestration of newly synthesized membrane proteins into ER whorls. Sequestration into ER whorls is exacerbated by HSP70 inhibition but can be rescued by overexpressing HSPA8 and HSPA1. Cross-linking assays reveal that 5’UTR binding sites stabilize eIF3–80S interactions during early elongation, facilitating recruitment of HSPA8 to ribosomes. These findings indicate that genetic instructions within 5’UTRs direct eIF3-mediated chaperone recruitment, ensuring proper co-translational folding of ER membrane proteins.

## Introduction

Cellular homeostasis is crucially dependent on the correct, co-translational folding of nascent peptides into functional proteins ^1–4^. Misfolded proteins can form toxic aggregates that cause various age-related diseases including cancer, neurodegeneration and metabolic disease ^5^. Co-translational folding is carried out by an elaborate network of chaperones strategically positioned in the vicinity of the ribosome ^2,3,6–8^. Among these chaperone complexes, the ribosome-associated complex (RAC) comprises a specialized HSP70 protein (HSPA14) and a cognate HSP40 J-domain protein (DNAJC2) ^9–11^. This assemblage appears to act on nascent chains in series with other HSP70 chaperones, such as SSB in yeast and HSPA1 and HSPA8 in mammalian cells ^11,12^. SSB-type chaperones facilitate co-translational protein folding, expedite protein synthesis, and promote efficient membrane targeting ^7,12–14^.

While yeast SSB chaperones are directly anchored to the ribosome ^15,16^, the mechanism by which mammalian HSP70s are recruited to nascent polypeptide chains remains less well understood. Our research has provided evidence that, for a subset of mRNAs, the translation initiation factor eIF3 may be involved in the recruitment of chaperones to the ribosome ^17,18^. eIF3 is a 13-subunit protein complex that plays various roles in canonical cap-dependent and non-canonical translation initiation ^19–22^. Despite eIF3’s global role in translation, loss-of-function phenotypes of individual eIF3 subunits are remarkably dissimilar, suggesting that some eIF3 subunits have mRNA-selective functions in translation. These findings can be reconciled by a model in which an ancestral eIF3a, b, g, i, j complex, conserved in budding yeast, performs basic functions in translation initiation, while additional subunits form modules with mRNA-selective functions ^17,23–25^.

Traditionally, eIF3 was thought to dissociate from ribosomes upon 60S subunit joining ^26,27^, but more recent evidence indicates that eIF3 can remain associated with 80S ribosomes during early elongation ^17,28–30^, particularly on mRNAs encoding membrane-associated proteins in higher eukaryotes ^17^. Depletion of eIF3e in non-transformed human MCF10A cells interferes with early translation elongation of approximately 2,700 mRNAs encoding proteins with membrane-associated functions ^17^. Under these conditions, several HSP70 chaperones are reduced in 80S ribosomal fractions ^17^, suggesting that eIF3e facilitates chaperone recruitment to ribosomes to coordinate translation initiation with co-translational protein folding.

A primary unresolved question of this model is how eIF3 senses at the start codon that it is translating an mRNA that will require its continued presence on the 80S ribosome downstream of the start codon to assist with nascent protein folding. We show here that this instruction is genetically programmed in mRNA 5’UTRs through distinct eIF3 binding sites. mRNA-assisted eIF3-80S tethering then promotes co-translational folding of ER membrane proteins via eIF3-mediated chaperone recruitment.

## Results

### eIF3 binding sites in 5’UTRs are required for proper localization of ER membrane-associated proteins

In previous work, we identified a set of ∼2,700 mRNAs that required eIF3e for efficient translation elongation within the first ∼75 codons; this set was highly enriched for membrane-associated functions including those of the endoplasmic reticulum (ER) ^17^. To determine the function of eIF3e in the expression of individual ER-associated proteins, we chose three mRNAs from our list of ∼2,700 eIF3e-dependent mRNAs and expressed them in MCF7 cells and AD293 cells as C-terminal enhanced green fluorescent protein (EGFP) fusions under control of their cognate 5’UTRs (**Figure 1A**). LRRC59 is a tail anchored ER membrane protein facing the cytosol, CANX a protein with a single transmembrane domain mostly extending into the ER lumen, and TMEM33 an integral ER and nuclear membrane protein with three transmembrane domains (**Figure 1B**). As indicated by PAR-CLIP data, each of these mRNAs contains bindings sites for eIF3 in its 5’UTR ^31,32^, which are predicted, based on SHAPE data ^33,34^, to form stable stem loop structures (**Figure 1C**).

**Figure 1.**
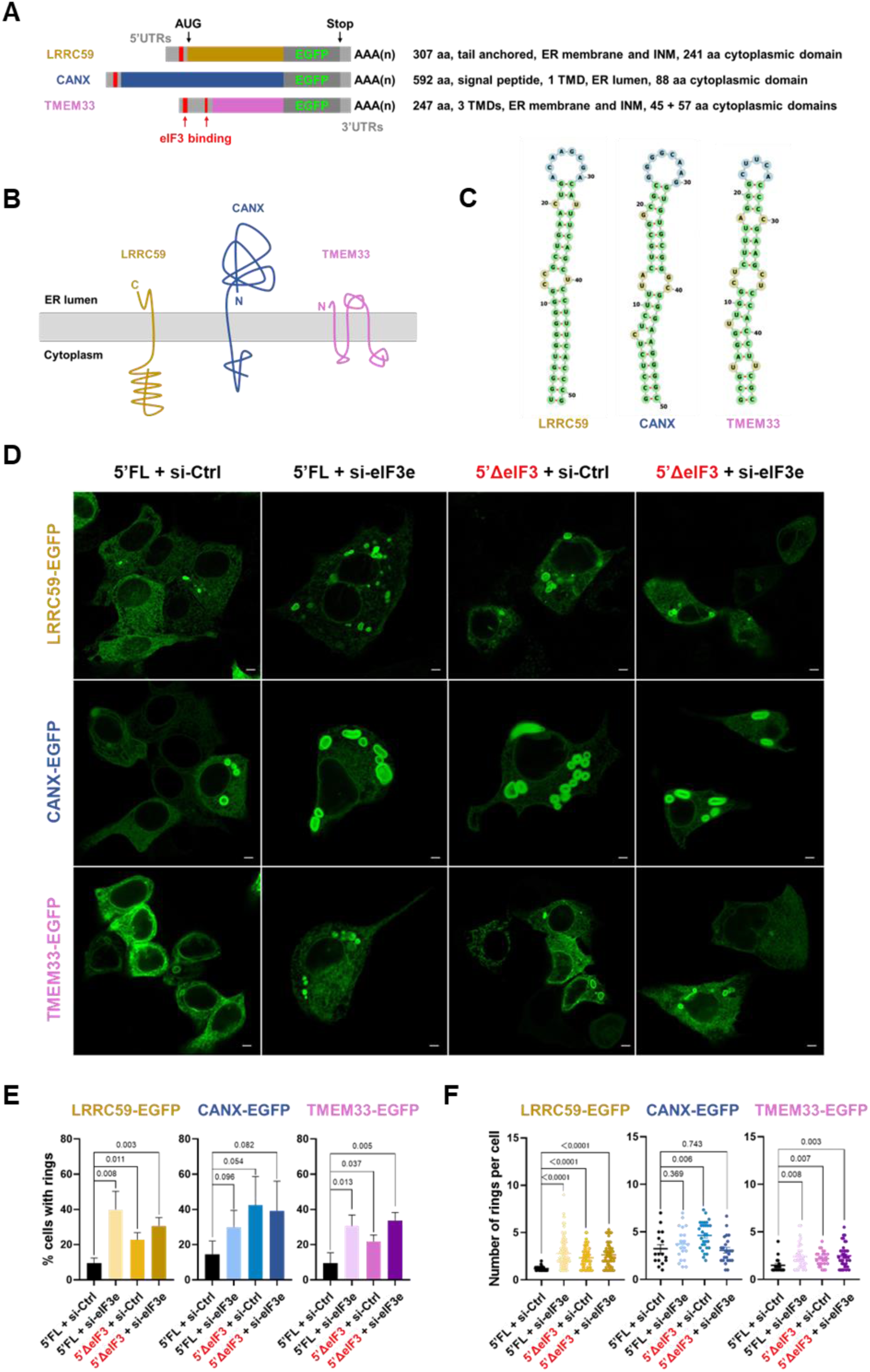
Effect of eIF3 on the localization of ER membrane proteins. **A.** Schematic of the mRNAs encoding ER membrane proteins fused to enhanced green fluorescent protein (EGFP) and expressed from plasmids. AAA(n) denotes poly-A tails, UTRs, untranslated regions. eIF3 binding sites in 5’UTRs mapped by CLIP are indicated in red. Protein features are summarized to the right. aa, amino acids, ER, endoplasmic reticulum; INM, inner nuclear membrane; TMD, transmembrane domain. **B.** Symbolic diagram of the topologies of the indicated ER membrane proteins. **C.** Secondary structures of the indicated mRNAs based on in vivo selective 2′-hydroxyl acylation analyzed by primer extension (SHAPE) data ^33,34^. LRRC59, ENST00000225972.8 Nucleotides116-166; CANX, ENST00000247461.9 Nucleotides 54-103; TMEM33, ENST00000264452.9 Nucleotides 42-87. **D.** The proteins shown in A. were expressed in MCF7 cells treated with control siRNA (si-Ctrl) or siRNA targeting eIF3e (si-eIF3e) for 72h. 24h after transfection, EGFP fusion proteins were detected by fluorescence microscopy of live cells. The left two panels show plasmids containing the full length 5’UTRs (5’FL), the right two panels show plasmids in which the eIF3 binding sites were removed from the 5’UTRs (5’ΔeIF3). Scale bars: 5 μm. **E.** Images as shown in D. were quantitatively scored for percentage of EGFP positive cells containing rings. Bars represent means (n = >200 cells) ± standard deviations, numbers represent p values (two-stage step-up method of Benjamini, Krieger, and Yekutieli). **F.** Images as shown in D. were quantitatively scored for the number of rings per EGFP positive cell. Data represent means (n = >16 cells) ± standard deviations, numbers represent p values (unpaired t-test).

Fluorescence microscopy of live cells revealed typical ER localization patterns for LRRC59-EGFP, CANX-EGFP, and TMEM33-EGFP (**Figure 1D**). Upon knockdown of eIF3e, all three EGFP fusion proteins were still readily expressed but partially relocalized into 2 - 9 cytoplasmic ring-like structures in up to 40% of EGFP positive cells (**Figure 1D, E, F**). Similar rings were observed in ∼10 - 15% of control knockdown cells, but these mostly occurred as singlets (**Figure 1D, E, F**). Knockdown of eIF3d, a subunit dimerizing and co-regulated with eIF3e ^24,35,36^, led to similar localization of LRRC59-EGFP in rings (**Figure S1A**), indicating that the integrity of the eIF3d/e module is required for proper subcellular targeting of this ER membrane protein. Sufficient eIF3 function appears to be retained in the knockdown cells to allow the synthesis of the EGFP-fused ER membrane proteins when overexpressed from plasmids.

To decipher the function of eIF3 in membrane targeting, we prepared similar plasmids in which the eIF3 binding sites in the 5’UTRs of *LRRC59, CANX*, and *TMEM33* were deleted (**Figure S1B,**). This manipulation of the non-coding region was sufficient to mislocalize the EGFP-fusion proteins into rings despite the eIF3 complex being intact (**Figure 1D - F**). Knockdown of eIF3e led to no or only a small additional increase in cytoplasmic rings, indicating that the effect of the 5’UTR binding site deletion was due to perturbing eIF3 activity acting on these mRNAs. Inverting the sequence of either one or both strands of the predicted stem loop sequence of the *LRRC59* mRNA also led to localization of LRRC59-EGFP into cytoplasmic rings (**Figure S1C**). This suggests that ring formation was not due to merely shortening of the 5’UTR but to disruption of an important secondary structure element bound by eIF3.

We subsequently employed CRSPR/Cas9 genome editing to delete eIF3 binding sites in the 5’UTRs of the endogenous *LRRC59*, *CANX*, and *TMEM33* genes in MCF7 and AD293 cells, the edited cell lines being referred to as 5’ΔeIF3 cell lines (**Figure 2A, S2A**). RNA immunoprecipitation using eIF3b antibodies revealed a ∼40 - 60% reduction in eIF3 binding to *LRRC59*, *CANX*, and *TMEM33* mRNAs containing the edited 5’UTRs lacking eIF3 binding sites thus confirming the published CLIP data ^31,32^ (**Figure 2B, S2B**). eIF3 binding to non-edited mRNA encoding *JUNB* was not reduced in any of the edited cell lines (**Figure 2B, S2B**). Deletion of eIF3 binding sites did not completely abolish eIF3-mRNA interactions, likely reflecting independent interactions mediated through the cap-binding complex.

**Figure 2.**
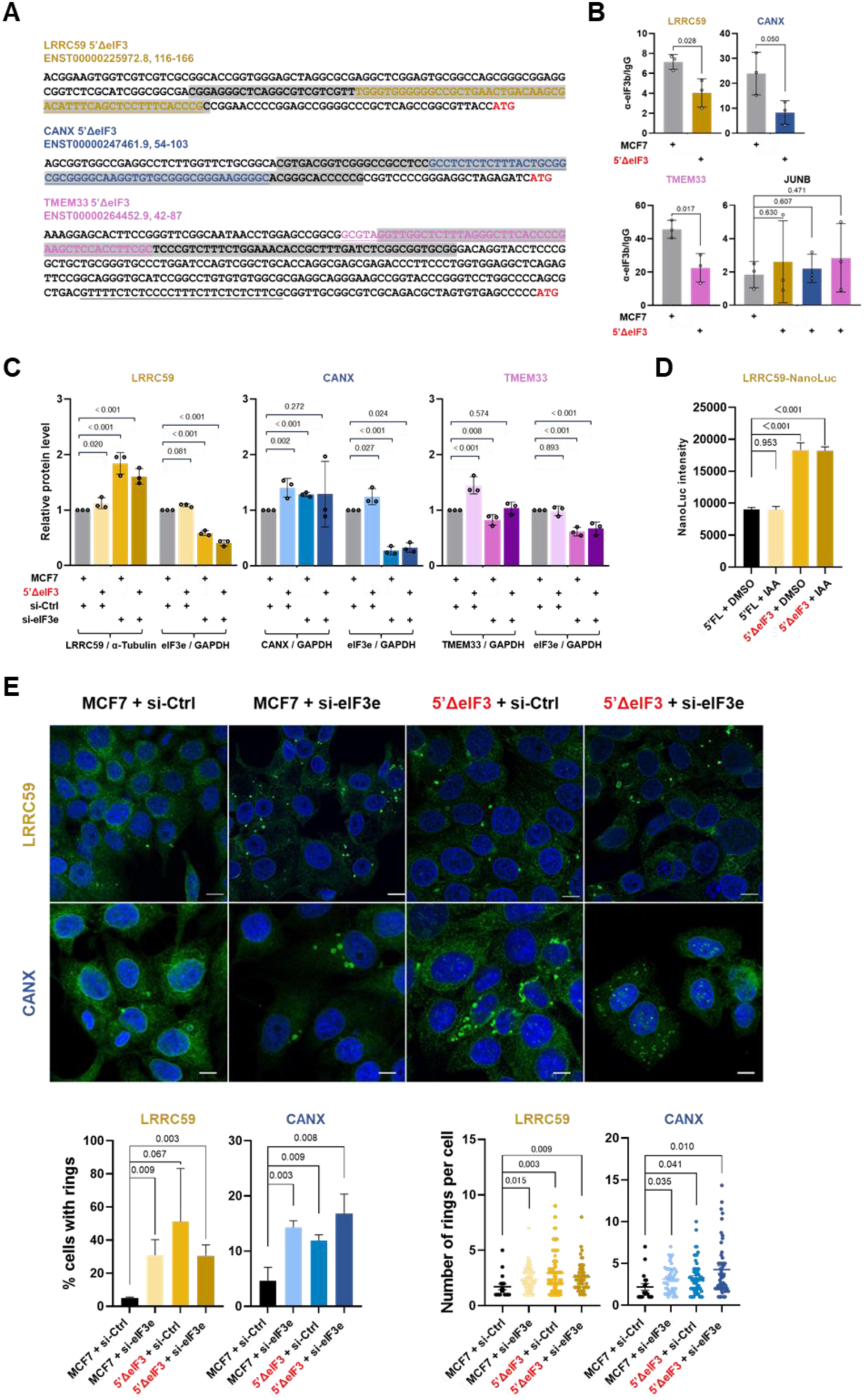
Effect of eIF3 5’UTR binding sites on the localization of ER membrane proteins. **A.** Sequences of the 5’UTRs of mRNAs encoding the indicated membrane proteins. The underlined sequences represent the eIF3 binding sites mapped by CLIP. The grey areas highlight the sequences deleted by genome editing in MCF7 cells. Start codons are highlighted in red font. The 5’UTR of *TMEM33* contains a second potential eIF3 binding site (underlined) which was considered minor based on the rarity of corresponding CLIP fragments and was therefore not edited. **B.** mRNPs present in cell lysate from parental MCF7 cells or from MCF7 cells in which the indicated eIF3 binding regions were deleted (5’ΔeIF3 cell lines) were immunopurified with eIF3b antibodies. Co-purification of the indicated mRNAs was quantified by RT-qPCR. Bars represent means (n = 3) ± standard deviations, numbers represent p values (two-stage step-up method of Benjamini, Krieger, and Yekutieli). **C.** The Western blots in Figure S2C were quantified to determine the effects of deleting eIF3 5’UTR binding sites (5’ΔeIF3) or knocking down eIF3e for 72 h on the steady-state levels of the indicated ER membrane proteins. The signals of ER membrane proteins were normalized to reference proteins (GAPDH or a-tubulin) as indicated. Bars represent means (n = 3) ± standard deviations, numbers represent p values (two-stage step-up method of Benjamini, Krieger, and Yekutieli). **D.** In vitro translation of nano-luciferase mRNA under the control of the *LRRC59* 5’UTR with or without the eIF3 binding sites (5’ΔeIF3). Translation competent cell lysate was prepared from HCT116 cells in which the endogenous copies of the eIF3e gene were modified with auxin-inducible degrons allowing rapid depletion of eIF3 by the addition of 500 μM indole-3-acetic acid (IAA) for 12 h (**Figure S2E**). Translation activity was determined by luciferase assay and confirmed by immunoblotting (**Figure S2E**). Bars represent means (n = 3) ± standard deviations, numbers represent p values (two-stage step-up method of Benjamini, Krieger, and Yekutieli). Western blots document depletion of eIF3e and synthesis of full-length nano-luciferase protein. **E.** Parental MCF7 cells or MCF7 cells in which the indicated eIF3 binding regions were deleted (5’ΔeIF3 cell lines) and/or eIF3e was knocked down for 72 h were fixed and stained with antibodies against LRRC59 or CANX. Bright ring-like structures were visualized by confocal microscopy, and the percentage of ring containing cells were scored (top panels). Bars represent means (n = >200 cells) ± standard deviations, numbers represent p values (two-stage step-up method of Benjamini, Krieger, and Yekutieli). In addition, the number of rings per cell was scored (bottom panels) with data representing means (n = >39 cells) ± standard deviations and numbers representing p values (unpaired t-test). Scale bars: 10 μm.

The expression levels of LRRC59, CANX, and TMEM33 proteins were not consistently reduced in 5’ΔeIF3 cell lines (**Figure 2C, S2C, D**). In addition, the eIF3 binding site of the *LRRC59* 5’UTR was dispensable for efficient translation in vitro as shown by a luciferase reporter assay with cell lysate of human HCT116 cells (**Figure 2D**). Indeed, deletion of the eIF3 binding site even enhanced in vitro translation of nano-luciferase under control of the *LRRC59* 5’UTR, possible due to removal of inhibitory secondary structure (**Figure 2D**).

In contrast, removing eIF3 binding sites from *LRRC59* and *CANX* mRNAs was sufficient to cause a ∼3 to 10-fold increase in cytoplasmic ring structures positive for LRRC59 and CANX proteins, respectively (**Figure 2E**). The same observations were made upon editing 5’UTRs in AD293 cells (**Figure S2E**). We were unable to assess the localization of endogenous TMEM33 protein because we were unsuccessful in sourcing or raising antisera that can detect TMEM33 by immunofluorescence staining (but see electron micrographs in **Figure 3A** below).

**Figure 3.**
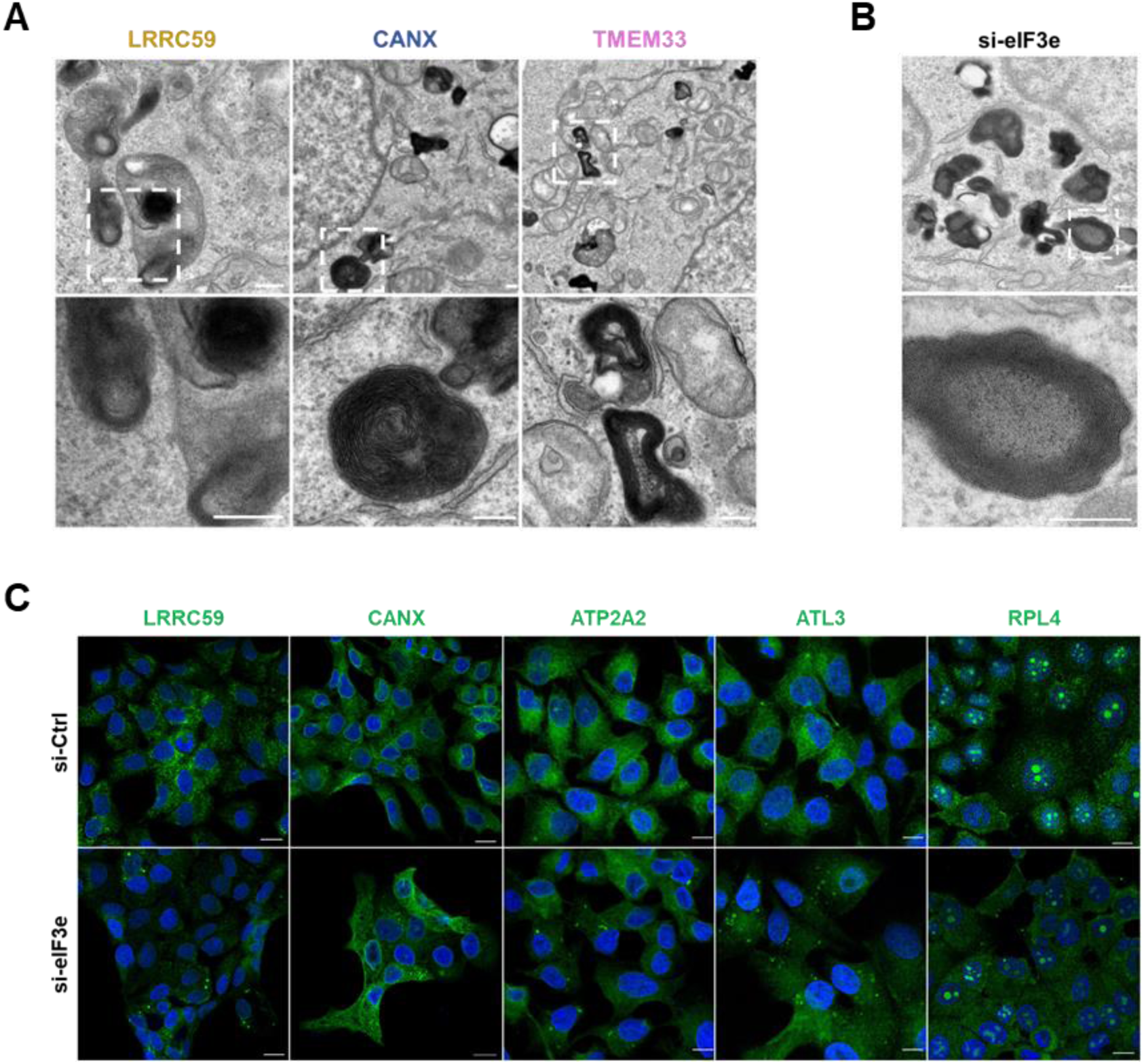
Formation of ER whorls in eIF3 deficient cells. **A.** MCF7 cells in which the eIF3 binding regions were deleted from the indicated genes (5’ΔeIF3 cell lines) were imaged by transmission electron microscopy. Dense membrane structures resembling ER whorls are shown. The bottom row shows magnifications of the region highlighted on the top row. Scale bars: 2 μm. **B.** Same as in A. with MCF7 cells in which eIF3e was knocked down with siRNA for 72 h. **C.** Immunofluorescence staining of the indicated proteins (green) in MCF7 cells transfected with control (si-Ctrl) or eIF3e targeting siRNA (si-eIF3e). RPL14 is localized in the cytoplasm and in nucleoli.

Knockdown of eIF3e in parental MCF7 and AD293 cells also caused accumulation of endogenous LRRC59 and CANX in cytoplasmic rings similar in extent to that seen with 5’ΔeIF3 cell lines (**Figure 2E, S2E**). However, knockdown of eIF3e in the edited cell lines did not cause further increases in the number of cytoplasmic rings (**Figure 2E, S2E**), again indicating that the 5’UTR binding site alterations perturb eIF3 action on these mRNAs. Overall, this data suggests that eIF3e is critical for directing proper ER membrane protein localization via mRNA 5’UTR binding sites, though not limiting for their expression.

### ER membrane proteins synthesized during eIF3 deficiency are misfolded and sequestered into ER whorls

The cytoplasmic rings we observed in eIF3e deficient cells resembled so-called ER whorls, representing multilayered rings of smooth ER that were linked to expression of mutant ER membrane proteins ^37,38^, ER stress ^39^, and protein misfolding ^40^. Electron microscopy revealed concentrically stacked membrane cisternae in 5’ΔeIF3 AD293 cells carrying edited 5’UTRs of *LRRC59*, *CANX*, and *TMEM33* (**Figure 3A**). Similar structures were detected in parental MCF7 cells upon knockdown of eIF3e (**Figure 3B**). The membrane stacks did not seem to be decorated with ribosomes and did not stain with antibodies against RPL14 (**Figure 3A-C**). However, they were strongly positive for ER membrane proteins (LRRC59, CANX, ATL3, and ATP2A2) by immunofluorescence staining (**Figure 3C**), indicating that they represent accumulations of smooth ER membranes similar to ER whorls ^37–40^. Consistent with this is the observation that formation of LRRC59 positive ER whorls induced by knockdown of eIF3e was mimicked by 600 nM of the ER stress inducer thapsigargin (TG) (**Figure S3A**). Withdrawal of TG for 6 hours reversed ER whorl formation as described ^39^, but this reversal was blocked by knocking down eIF3e (**Figure S3A**). However, unlike TG, neither knockdown of eIF3e nor deletion of eIF3 binding sites in 5’UTRs induced any signs of ER stress as determined by eIF2α phosphorylation, ATF4 and BiP (HSPA5) accumulation, or *XBP1* mRNA splicing (**Figure S3B, C**). Thus, whereas ER stress can readily induce ER whorls ^39^, ER whorls resulting from eIF3 deficiency do not coincide with global, unresolved ER stress.

Our data suggested that ER membrane proteins synthesized with low levels of eIF3e or from mRNAs with reduced eIF3 binding in their 5’UTRs are trapped in ER whorls. To study this, we performed fluorescence reactivation after photobleaching (FRAP) studies. LRRC59, CANX, and TMEM33 fusions with EGFP were expressed in MCF7 cells from plasmids lacking eIF3 binding sites in their cognate 5’UTRs (**Figure S1B**). Fusion proteins located to areas of reticular ER or to ER whorls were photobleached, and fluorescence recovery was monitored over time. Whereas all three membrane proteins repopulated the bleached areas of the reticular ER within ∼1 - 2 minutes, essentially no fluorescence recovery was observed for ER whorls (**Figure 4A-C, Supplementary Movies 1 - 3**). This suggests that ER membrane proteins are stably compartmentalized to ER whorls when eIF3 function is compromised.

**Figure 4.**
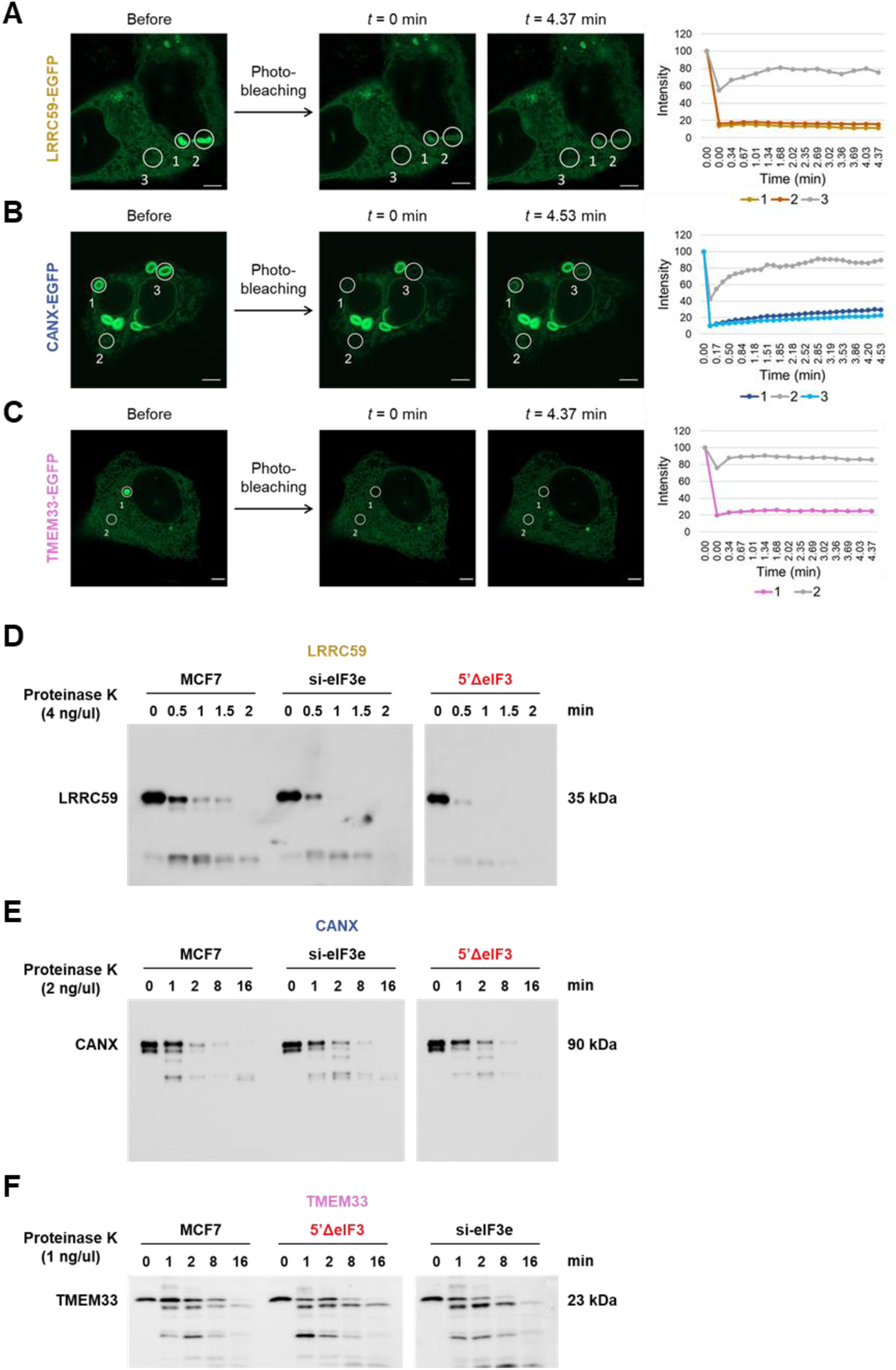
Effect of eIF3 on the folding state of ER membrane proteins. **A. - C.** MCF7 cells were transfected with plasmids encoding the indicated EGFP fusion proteins under control of their cognate 5’UTRs from which the eIF3 binding sites were deleted (5’ΔeIF3, **Figure S1B**). After 24 h, live cells were photobleached in the indicated areas of tubular ER or ER whorls, and relative fluorescence recovery was quantified over time (right panels). Scale bars: 5 μm. **D. – E.** The indicated ER membrane proteins were extracted from parental MCF7 cells or from MCF7 cells in which the indicated eIF3 binding regions were deleted (5’ΔeIF3 cell lines) or eIF3e was knocked down for 72 h. Extracts were subjected to limited proteolysis with proteinase K, and proteolytic products were detected by immunoblotting.

Previous studies have demonstrated sequestration into ER whorls of ER membrane proteins that were partially misfolded due to single residue disease variants ^37,38^. To assess the conformational maturation and folding state of ER membrane proteins synthesized with compromised eIF3 function, we performed limited proteolysis of LRRC59, CANX, and TMEM33 upon extraction from the membrane fraction. ER membrane proteins synthesized in eIF3e knockdown cells or in UTR-edited 5’ΔeIF3 cell lines were substantially more sensitive to digestion with proteinase K (**Figure 4D-F, S4A**), indicating that they are not in their native conformations. The increases in proteinase K sensitivity were selective for the respective proteins whose mRNA 5’UTRs were edited. For example, LRRC59 was only protease sensitive in LRRC59 5’ΔeIF3 cells, not in CANX or TMEM33 5’ΔeIF3 cell lines and vice versa (**Figure S4B**). Thus, binding of a functional eIF3 complex to mRNA 5’UTRs is required for efficient folding of newly synthesized ER membrane proteins.

### eIF3 promotes chaperone recruitment to the ribosome

Our previous proteomic data in yeast and mammalian cells revealed interactions of eIF3 with 80S ribosomes and several HSP70 chaperones as well as the CCT/TRiC chaperonin complex ^17,18^ all of which are known to be involved in co-translational protein folding ^41–43^. We found that HSPA1 and HSPA8 are the most abundant HSP70 chaperones co-fractionating with eIF3-80S complexes in MCF10A and HeLa cells ^17^. We therefore assessed the effect of the HSP70 inhibitor VER-155008, which equally inhibits HSPA1 and HSPA8 ^44^, on the partitioning of LRRC59 into ER whorls. VER-155008 led to a pronounced increase in LRRC59 whorls, similar in extent to knockdown of eIF3e (**Figure 5A**, compare 1E, F). Similarly, knockdown of HSPA1 and HSPA8 led to LRRC59 positive ER whorl formation (**Figure S5A**). Additionally, knockdown of CCT/TriC subunits CCT1 and CCT8 caused ER whorls (**Figure 5C**). The data suggested that eIF3-80S-associated chaperone activity may be required for proper folding of LRRC59.

**Figure 5.**
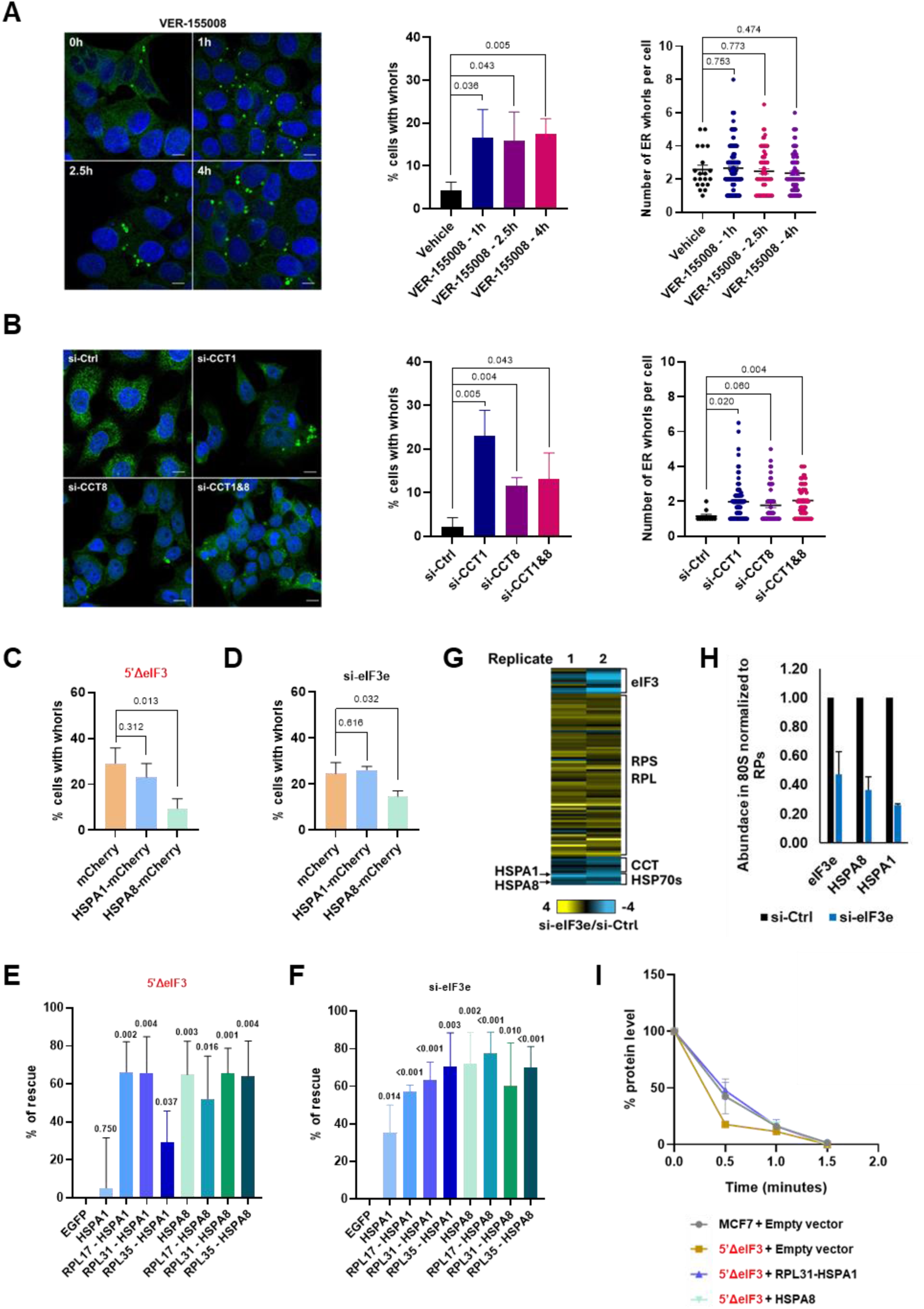
Effect of eIF3 on chaperone-mediated folding of LRRC59. **A.** MCF7 cells were treated with the broad spectrum HSP70 inhibitor VER-155008 for the indicated periods of time. Cells were fixed and LRRC59 positive ER whorls were detected by indirect immunofluorescence staining. The percentage of cells containing ER whorls was scored (left panel). Bars represent means (n = >200 cells) ± standard deviations, numbers represent p values (two-stage step-up method of Benjamini, Krieger, and Yekutieli). In addition, the number of ER whorls per cell was scored (right panel) with data representing means (n = >44 cells) ± standard deviations and numbers representing p values (unpaired t-test). Scale bars: 10 μm. **B.** Same as in B. but knockdown was for the indicated subunits of the CCT/TRiC chaperonin complex. Scoring was done as in A (n > 15 cells). **C., D.** MCF7 cells in which the eIF3 binding region was deleted from the 5’UTR of *LRRC59* (C.) or that were transfected with si-eIF3e for 72 h (D.), were transfected with plasmids driving the expression of mCherry or mCherry fused to the C-termini of HSPA1 or HSPA8. The percentage of cells showing LRRC59 positive ER whorls were scored. Bars represent means (n = >200 cells) ± standard deviations, numbers represent p values (two-stage step-up method of Benjamini, Krieger, and Yekutieli). **E., F.** Same experiment as in C., D. for cells transfected with plasmids driving the expression of EGFP (negative control) and HSP70 proteins either unfused or fused to the indicated 60S ribosomal proteins at their N-termini. LRRC59 positive ER whorls were scored, and results are displayed as percent reduction in ER whorl containing cells relative to the EGFP transfected samples (% rescue). Bars represent average percent rescue (n = 3) ± standard deviations. Numbers represent p values (two-stage step-up method of Benjamini, Krieger, and Yekutieli). **G.** Cell lysates from MCF7 cells transfected with control (si-Ctrl) or eIF3e targeting siRNA (si-eIF3e) were separated by sucrose density gradient centrifugation. Fractions containing 80S ribosomes were analyzed by quantitative LC-MS/MS. Protein ratios (si-eIF3e/si-Ctrl) of replicate experiments are shown in a heatmap (left). RPS and RPL indicate 40S and 60S ribosomal proteins, respectively. **H.** Relative abundance of eIF3e and chaperones in 80S fractions of si-Ctrl and si-eIF3e transfected cells normalized to the sum abundance of all detected ribosomal proteins; see Data File S1. **I.** MCF7 cells in which the eIF3 binding region was deleted from the 5’UTR of *LRRC59* were transfected with plasmids driving the expression of HSPA8 fused to mCherry or HSPA1 fused to RPL31. LRRC59 was extracted and subjected to limited proteolysis with proteinase K, and proteolytic products were detected by immunoblotting (see Figure S5J).

To test this, we asked whether overexpression of HSPA1 and HSPA8 could rescue the formation of LRRC59 positive ER whorls resulting from eIF3 deficiency. mCherry-HSPA8 reduced ER whorl formation by ∼40 - 65% (**Figure 5D, E; S5B, C**). mCherry-HSPA1 overexpression was less efficient; it reduced ER whorls by ∼20% in cells in which the eIF3 binding site was deleted from the 5’UTR of *LRRC59* mRNA (**Figure 5D; S5B, C**) but failed to rescue ER whorl formation in eIF3e knockdown cells (**Figure 5E; S5B, C**).

Given the more intimate interaction of constitutive HSPA8 with ribosome-nascent chain complexes compared to heat-inducible HSPA1 ^11^, we tested whether forced recruitment of HSPA chaperones to ribosomes would improve their rescue activity. N-termini of both chaperones were fused to 60S ribosomal proteins surrounding the ribosome exit tunnel. Among these were RPL35/uL29 known to interact with SSB-type HSP70 chaperones in yeast, RPL31/eL31 which interacts with RAC ^45,46^, and RPL17/uL22 which neighbors RPL31 (**Figure S5D, E**). Fusion with all three RPLs greatly augmented the ability of HSPA1 to rescue LRRC59 positive ER whorl formation in the presence of eIF3 deficiency (**Figure 5F, G; S5F, G**). In contrast, fusion with RPL proteins did not affect rescue by HSPA8 (**Figure 5F, G; S5F, G**). This data suggests that HSPA1 can efficiently rescue eIF3 deficiency in ER membrane protein folding when ectopically anchored to a position at the ribosome that is equivalent to SSB in yeast ^15,16^, whereas HSPA8 does not appear to require anchoring.

This data suggested a role of eIF3 in the recruitment of HSP70 chaperones to 80S ribosomes. To determine this, we performed quantitative proteomics on 80S monosomes isolated from MCF7 cells by sucrose density gradient fractionation. Knockdown of eIF3e led to the typical changes in the composition of holo-eIF3 present in 80S fractions with depletion of all subunits except the ancestral core complex consisting of eIF3a, b, g, i, and j ^17,23–25^ (**Figure S5H, Data File S1**). In addition, knockdown of eIF3e caused an increase in the 80S peak (**Figure S5I**) with a corresponding increase in 40S and 60S ribosomal proteins ribosomal proteins as determined by LC-MS/MS of 80S fractions (**Figure 5H**). In contrast, levels of HSP70 and CCT/TRiC chaperones were reduced in 80S fractions isolated from eIF3e knockdown cells (**Figure 5H, Data File S1**). Considering the increase in 80S peak size, eIF3e knockdown led to a 63 – 74% reduction of HSPA8 and HSPA1 after normalization to ribosomal protein abundance (**Figure 5I, Data File S1**). Conversely, HSP70 proteins rescued the misfolding of LRRC59 resulting from eIF3 deficiency. We observed that proteinase K hypersensitivity of LRRC59 protein in LRRC59 5’ΔeIF3 cells was efficiently rescued by overexpressing HSPA8 or RPL31-HSPA1 (**Figure 5J, S5J**), indicating that high levels of both chaperones can mitigate LRRC59 misfolding resulting from eIF3 deficiency, presumably due to ectopic recruitment to ribosomes. Taken together, these data suggest that eIF3 promotes recruitment of chaperones to 80S ribosomes, thereby ensuring correct folding of ER membrane proteins during synthesis.

### eIF3 binding sites in 5’UTRs stabilize eIF3-80S interactions

To fulfill such a function, eIF3 would need to stay on the 80S ribosome until nascent chains emerge from the ribosome exit tunnel, typically after 30 – 60 codons. We adapted a cross-linking method ^47^ to measure the abundance of eIF3-80S complexes in the start codon proximal regions of *LRRC59*, *CANX*, and *TMEM33* mRNAs (**Figure 6A**). MCF7 cells were first subjected to cross-linking with dithiobis(succinimidyl propionate), mRNA-protein complexes were isolated, and start codon proximal mRNA regions were efficiently excised by oligonucleotide directed cleavage with RNAse H (**Figure S6A, B**). Samples were further separated by sucrose density gradient centrifugation, and 80S containing fractions (including polysomes) were collected. eIF3-80S complexes were immunoprecipitated with anti-eIF3b antibodies and attached *LRRC59*, *CANX*, and *TMEM33* mRNA fragments were quantified by qRT-PCR. Rabbit IgG immunoprecipitates were used as specificity controls.

**Figure 6.**
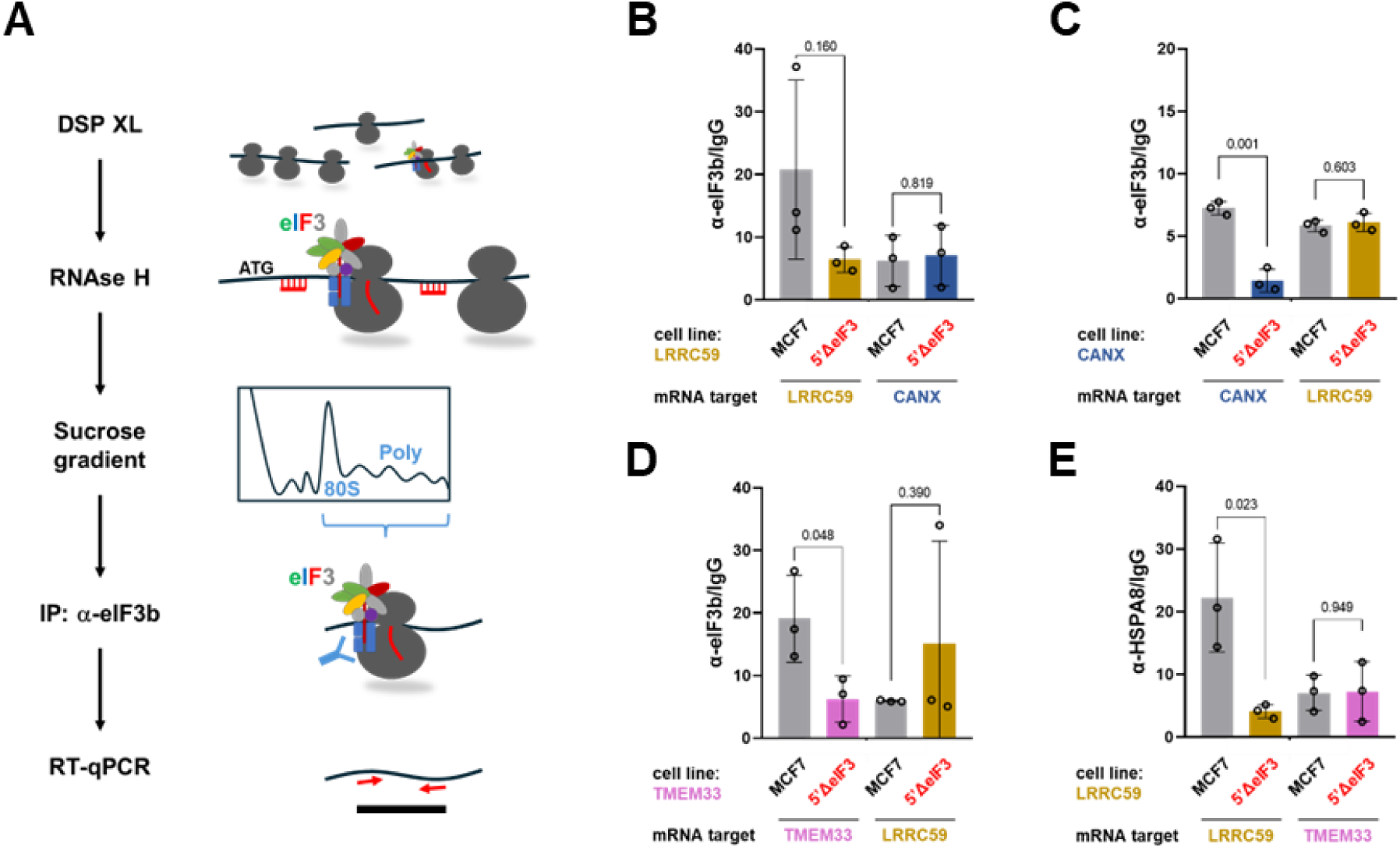
Effect of 5’UTR eIF3 binding sites on the stability of start codon proximal eIF3-80S. **A.** Schematic of the cross-linking protocol. MCF7 cells were first subjected to cross-linking with dithiobis(succinimidyl propionate) (DSP XL), mRNA-protein complexes were isolated, and start codon proximal mRNA regions were excised by oligonucleotide directed cleavage with RNAse H (oligos indicated in red, see Figure S6A). Samples were subsequently separated by sucrose density gradient centrifugation, and 80S containing fractions including polysomes were collected. eIF3-80S complexes were immunoprecipitated with anti-eIF3b or anti-HSPA8 antibodies and attached *LRRC59*, *CANX*, and *TMEM33* mRNA fragments were quantified by RT-qPCR with gene specific primers (indicated as red arrows). **B.** Cross-linking assay to measure the abundance of eIF3-80S complexes between codons 24 - 107 of *LRRC59* mRNA. Lysate from parental MCF7 cells and MCF7 cells in which the eIF3 binding region was deleted from the 5’UTR of *LRRC59* were used for anti-eIF3b or IgG (negative control) IP. The retained mRNA fragments were detected by RT-qPCR with primers specific for *LRRC59* (2 bars on the left) or *CANX* mRNA (specificity control, 2 bars on the right). Bars represent average enrichment of mRNA in anti-eIF3b IPs relative to IgG (n = 3, ± standard deviations); numbers represent p values (two-stage step-up method of Benjamini, Krieger, and Yekutieli). **C.** Cross-linking assay to measure the abundance of eIF3-80S complexes between codons 19 - 119 of *CANX* mRNA. Same experiment as in B. but with MCF7 cells in which the eIF3 binding region was deleted from the 5’UTR of *CANX. LRRC59* was used as a specificity control. Bar graph as in B. **D.** Cross-linking assay to measure the abundance of eIF3-80S complexes between codons 32 - 76 of *TMEM33* mRNA. Same experiment as in B. but with MCF7 cells in which the eIF3 binding region was deleted from the 5’UTR of *TMEM33. LRRC59* was used as a specificity control. Bar graph as in B.

The experiments revealed binding of eIF3 to 80S complexes within the first ∼20 – 100 codons of *LRRC59, CANX*, and *TMEM33* mRNA (**Figure 6C-E**). These findings confirmed our previous eIF3b and eIF3e selective ribosome profiling data ^17^. Deletion of 5’UTR eIF3 binding sites in 5’ΔeIF3 cell lines led to substantially reduced retention of eIF3 on post-initiation 80S ribosomes (**Figure 6C-E**). The reductions were selective for the respective mRNA whose 5’UTRs were edited. For example, deletion of the eIF3 binding site in *LRRC59* mRNA affected eIF3-80S interaction only on *LRRC59* mRNA but not on *CANX* and *TMEM33* mRNAs (**Figure 6C**). Similar data was obtained with 5’UTR edited AD293 cells (**Figure S6C - E**).

This data strongly suggested that eIF3 binding sites in the 5’UTRs of *LRRC59, CANX*, and *TMEM33* mRNAs stabilize eIF3-80S interactions during early translation elongation, likely to promote chaperone recruitment to nascent ER membrane proteins. To test this conjecture, we applied the cross-linking assay to HSPA8. Consistent with reduced formation of eIF3-80S complexes, cells in which eIF3 binding sites in the 5’UTRs of *LRRC59* mRNA were deleted showed reduced recruitment of HSPA8 to 80S ribosomes during early translation elongation (**Figure 6F**).

## Discussion

Our findings support a model of eIF3 function wherein this factor coordinates translation initiation with chaperone recruitment to ribosomes thereby facilitating the timely folding of nascent ER membrane proteins (**Figure 7**). These results align with our previous observations that (i) eIF3 forms stable complexes with 80S ribosomes, translation elongation factors, and chaperones ^18^, and (ii) eIF3 is critical for the early elongation of mRNAs encoding membrane-associated proteins ^17^. Specifically, we have identified a critical role for eIF3 in promoting the recruitment of HSPA8/1 and the CCT/TriC chaperonin to ribosomes, which is crucial for the correct folding of three ER membrane proteins with distinct topologies. This underscores the overarching importance of eIF3 in the accurate synthesis of ER membrane proteins.

**Figure 7.**
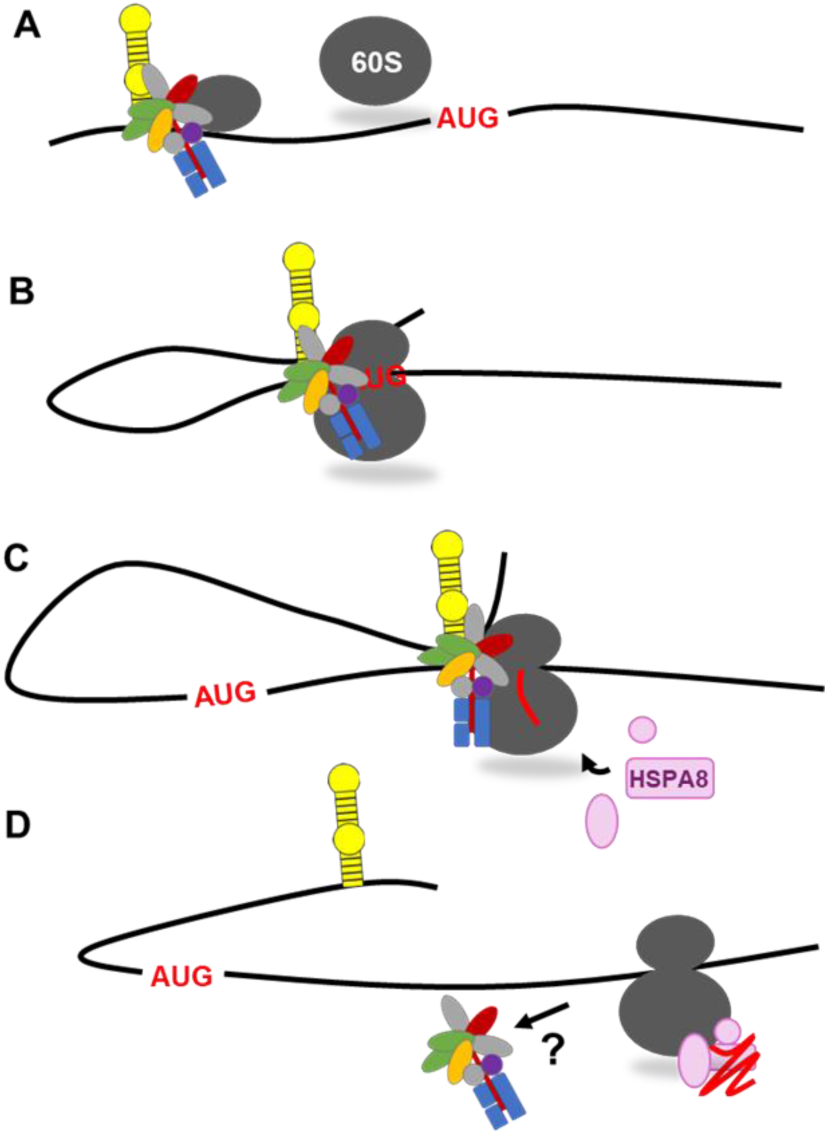
Model for 5’UTR-assisted stabilization of the eIF3-80S complex for chaperone recruitment. **A.** The eIF3-40S complex together with other initiation factors (not shown) binds to the mRNA. Before or during scanning, eIF3 interacts independently with a secondary structure element (depicted as hairpin) of the 5’UTR. **B.** eIF3-5’UTR interaction is maintained upon start codon recognition and subunit joining, forming the eIF3-80S complex. **C.** eIF3-80S stabilized through eIF3-5’UTR interactions is retained on the mRNA during early elongation to recruit HSPA8 and other chaperones. **D.** Once chaperone recruitment to nascent membrane proteins has occurred, eIF3 dissociates from 80S complexes through an unknown mechanism.

### Instructions for membrane protein folding are encoded in the non-coding genome

To perform its post-initiation function, eIF3 must remain associated with 80S ribosomes following the joining of 40S and 60S subunits. Once considered implausible due to steric hindrance, recent structural studies have strongly suggested that eIF3 can indeed be retained on 80S ribosomes ^48^. In addition, selective ribosome profiling has demonstrated interaction of eIF3 with active 80S ribosomes ^17,28–30^. Across the global translatome, eIF3 dissociates from 80S ribosomes stochastically with a decay half-length of ∼13 codons ^28^. Dissociation is substantially delayed to 100 – 150 codons in a subset of mRNAs enriched in membrane-associated functions, including *CANX* and *TMEM33* (*LRRC59* was not detected in that study) ^17^. Our data indicate that the signal prompting eIF3 to extend its stay on the 80S ribosome resides in eIF3 binding sites within the mRNA 5’ UTRs, not within the nascent proteins. Deletion of these binding sites diminishes the stability of the eIF3-80S complex on mRNA and reduces HSPA8/1 recruitment, leading to ER membrane protein misfolding and their sequestration into ER whorls. Thus, essential cues for the efficient co-translational folding of ER membrane proteins are hard-wired into the genome, a finding carrying several important implications.

Firstly, this finding broadens the functional versatility of mRNA 5’UTRs. While 5’UTRs are known to influence translation initiation and efficiency, their role in directing co-translational folding is unprecedented. Few similar yet distinct scenarios have been reported for 3’UTRs, such as the pre-recruitment of the signal recognition particle to ribosomes for membrane targeting ^49^ and the use of 3’UTR-encoded genetic information to enable functional protein-protein interactions in TIGER domains ^50^. Secondly, our results raise the intriguing possibility that non-coding variants disrupting eIF3 mRNA binding sites, likely through alterations of their hairpin structures, may underlie human genetic disorders characterized by protein misfolding. Indeed, single nucleotide variants in eIF3 mRNA binding sites have previously been implicated in hereditary hyperferritinemia cataract syndrome ^51^.

### eIF3-mRNA interactions

Throughout this study, we disabled eIF3 function by knocking down the eIF3e subunit. Notably, knocking down eIF3e had no measurable effect on the total protein levels of LRRC59, CANX, and TMEM33. Similarly, deleting eIF3 binding sites in mRNAs encoding these ER membrane proteins did not reduce their expression or impair in vitro translation of a reporter mRNA under the control of the *LRRC59* 5’UTR. This suggests that (i) a minimal core complex consisting of eIF3a, b, g, i and j, which remains largely intact upon eIF3e knockdown ^17,23–25^, is sufficient for translation initiation on these mRNAs, and (ii) the 5’UTR eIF3 binding sites are not required for basal translation of these mRNAs in vivo. Consequently, the folding-promoting function of eIF3 described herein is likely mediated by eIF3e and its associated proteins, particularly including its dimerization partner, the alternative cap-binding protein eIF3d ^52^. However, the specific eIF3 subunits involved in chaperone recruitment remain unclear. Notably, budding yeast which lacks eIF3d and eIF3e subunits, may circumvent the need for eIF3 in chaperone recruitment through direct anchoring of SSB-type HSP70 to 60S ribosomal proteins and rRNA ^15,16^.

The mechanism by which eIF3 binds to 5’UTRs is not fully understood. Given the lack of apparent homology in the primary sequences of eIF3 5’UTR binding sites determined by CLIP ^31^, eIF3 mRNA binding most likely depends on secondary structure. Indeed, mutagenesis data suggest that the eIF3 binding elements identified in *ALKBH5* and *JUN* mRNAs form stem-loop structures ^31,53^, a finding confirmed by nuclear magnetic resonance structure determination of the motif in *JUN* mRNA ^54^. The three eIF3 binding sites we deleted from the 5’UTRs of *LRRC59, CANX*, and *TMEM33* also appear to fold into stem loops based on SHAPE data ^33,34^. It is thus conceivable that eIF3 interacts with these mRNAs in a manner similar to that observed for *JUN*.

In our model, the critical function of eIF3-5’UTR interaction is to stabilize the complex between eIF3 and 80S ribosomes during early elongation and chaperone recruitment (**Figure 7**). How eIF3 is eventually removed from 80S ribosomes remains unknown. Possible mechanisms include post-translational modification of eIF3 or the ribosome, or steric constraints that limit the size of the mRNA looped out between the eIF3 binding element and the downstream open reading frame (ORF). Increasing tension in the looped-out mRNA may eventually disrupt the eIF3-80S complex (**Figure 7D**).

### ER whorl formation as a quality control mechanism

Our data further suggest that sequestration into ER whorls serves as a quality control mechanism for misfolded ER membrane proteins. Several membrane proteins carrying disease-causing single residue variants, including PMP22 (p.Leu16Pro, causing Charcot-Marie-Tooth neuropathy) and torsin A (TOR1A p.Glu303del, causing early-onset torsion dystonia, DYT1), accumulate in ER whorls ^37,38^, where they may be cleared by ER phagy. How cells detect the presence of misfolded membrane proteins to initiate ER whorl formation remains unclear. ER whorl formation in response to ER stress induced by thapsigargin involves the COPII vesicle fusion machinery in concert with PERK activation ^39^. Notably, eIF3 deficiency did not lead to the induction of ER stress as measured by PERK activity, IRE1 splicing, or BiP induction. This suggests that eIF3 deficiency does not cause major protein misfolding stress within the ER lumen. Instead, the misfolding occurring at the cytosolic ER membrane appears to be resolved through the sequestration of defective proteins into ER whorls.

### Limitations of the study

As the present study was done in cancer cells cultured in vitro, the physiological significance of eIF3-dependent chaperone recruitment for protein folding remains unproven in a living organism. Secondly, we have studied a very limited set of three ER membrane proteins, albeit in two different cell lines, MCF7 and AD293. Future studies could resolve these issues, for example, by globally profiling protein conformation in tissues derived from conditional eIF3e knockout mice. A more mechanistic understanding will also require in vitro biochemical studies, including structural studies of the eIF3-80S complex which has remained elusive. It is possible that addition of an mRNA containing an eIF3 binding site may stabilize the complex sufficiently for high resolution structure determination. Finally, whereas our study suggests ER whorl formation as a potential protein quality control pathway, the sensors, signal transducers, and effectors participating in the sorting of misfolded ER membrane proteins into ER whorls will remain to be discovered.

## Acknowledgements

The support from the Equipment Platform of the State Key Lab of Cellular Stress Biology at Xiamen University and Westlake Laboratory of Life Sciences and Biomedicine is gratefully acknowledged. This work was partially funded through grant grants 81773771 and 31770813 from the National Science Foundation of China (D.A.W), the Natural Science Foundation of Fujian Province (2018J01053), the Fujian Province Science and Technology Pilot Project (2021Y0002), the Fundamental Research Funds for the Central Universities (20720220053), and the Innovation Program of Xiamen University Department of Life Sciences & Human Health (Y.C.). Later stages of the work were supported by Westlake Laboratory of Life Sciences and Biomedicine and the Pioneer and “Leading Goose” R&D Program of Zhejiang Province (2024SSYS0029).

## Author contributions

Conceptualization: B.H., D.A.W.; Methodology: B.H., S.Z., H.D., Y.Z., B.Y., W.L.; Formal Analysis: B.H., D.A.W., Investigation: B.H., S.Z., H.D., Y.Z., B.Y., W.L.; Writing – Original Draft: D.A.W.; Writing – Review & Editing: B.H., S.Z., H.D., Y.Z., B.Y., W.L., Y.C., D.A.W.; Visualization: B.H., D.A.W.; Supervision: Y.C., D.A.W.; Funding Acquisition: Y.C., D.A.W.

## Declaration of generative AI and AI-assisted technologies in the writing process

During the preparation of this work, the author(s) used WestlakeChat in order to maximize the readability to a wide audience. After using this tool or service, the author(s) reviewed and edited the content as needed and take(s) full responsibility for the content of the publication.

## Methods

### Cell lines

MCF7 (female) and AD293 (female) cells were cultured in complete growth medium consisting of DMEM/High Glucose (HyClone, SH30022.01), 10% fetal bovine serum, 100 units/ml penicillin, and 100 μg/ml streptomycin under a humified environment with 5% CO_2_ at 37 °C. The male HCT116 cell line expressing OsTIR1 ^55^ was obtained from the Riken Bioresource Research Center of Japan and was cultured in McCoy’s 5A medium (Thermo Fisher Scientific) supplemented with 10% FBS (Gibco), 2 mM L-glutamine, 100 units/ml penicillin, and 100 mg/ml streptomycin at 37 °C and 5% CO_2_. Cells were authenticated by short tandem repeat sequencing and periodically determined to be free of mycoplasma.

### Plasmids and cloning

eIF3 binding sites in 5’UTRs were identified from published datasets using the Postar3 platform ^56^. LRRC59, CANX and TMEM33 containing their cognate 5’UTRs were inserted into pcDNA3.0 vector and fused with EGFP at the C-terminus. The subsequent mutations in the *LRRC59*, *CANX* and *TMEM33* 5’-UTRs were generated using these initial constructs. For LRRC59-5’ΔeIF3 constructs, nucleotides 98–136 and 153–176 in the 5’-UTR of *LRRC59* mRNA were deleted or inverted using Q5® Site-Directed Mutagenesis Kit (NEB, E0552S) following the manufacturer’s instructions. For CANX-5’ΔeIF3 constructs, nucleotides 45–68 in the 5’-UTR of *CANX* mRNA were deleted. For TMEM33-5’ΔeIF3 constructs, nucleotides 28–79 and 153–176 in the 5’-UTR of LRRC59 mRNA were deleted. See Figure S1B for sequences.

### Knockdown of eIF3e

MCF7 and AD293 cells were seeded and grown for 24 hours to a density of ∼20 %. 20 nM control or eIF3e si-RNA were transfected using Lipofectamine 2000 for 72 h before harvesting cells for protein analysis. The si-RNA oligo sequences were:

si-control: 5 ʹ -UUCUCCGAACGUGUCACGUdTdT-3 ʹ;
si-eIF3e: 5 ʹ-AAGCUGGCCUCUGAAAUCUUAdTdT-3 ʹ ^36^.

### Immunoblotting

Cells were collected by scraping into 0.1 mL of SDS sample buffer (60 mM Tris/Cl, pH 6.8, 5% beta-mercaptoethanol, 2% SDS, 10% glycerol, 0.02% bromophenol blue) followed by heating samples for 10 minutes at 95 °C. Protein lysates were subjected to SDS-PAGE and transferred to PVDF membranes. The membranes were blocked with 5% nonfat powdered milk for 1 hour at room temperature and incubated with primary antibodies overnight at 4 °C. After washing the membranes three times with TBST (10 mM Tris/Cl, pH 7.6, 150 mM NaCl, 1% Tween 20) for 10 minutes each, they were incubated with goat anti-mouse or anti-rabbit secondary antibodies conjugated to horseradish peroxidase for 1 hour at room temperature. Membranes were washed three times as above, followed by chemiluminescence detection using ECL reagents. Blots that were used for quantifications were visualized with the ChemiDoc™ imaging system (BIO-RAD) in an exposure time accumulation approach, and the appropriate exposure signal were selected for quantification. Quantifications were performed using ImageJ software. Primary antibodies used were: Rabbit anti-LRRC59 (GeneTex, cat. GTX122902,), rabbit anti-CANX (Proteintech, cat. 10427-2-AP), rabbit anti-TMEM33 (Bethyl, cat. A305-597A), anti-eIF3b (Bethyl, cat. A301-761A), anti-eIF3e (Bethyl, cat. A302-985A).

### Immunofluorescence staining

Cells were grown on cover glasses (NEST, cat. 801008) and fixed with 4% paraformaldehyde (PFA) at room temperature for 15 min, then permeabilized with 0.1% Triton X-100 in PBS buffer for 5 min at 4 °C and coverslips were blocked with 10% bovine serum albumin (BSA) in PBS buffer at room temperature for 1 h. Samples were incubated with antibody in 10% BSA in PBS buffer at 4 °C overnight. The primary antibody used was rabbit anti-LRRC59 (GeneTex, GTX122902, 1:100), rabbit anti-CANX (Proteintech, 10427-2-AP, 1:100), mouse anti-SERCA2 ATPase (Novus Biologicals, NB300-581, 1:100), rabbit anti-ATL3 (Proteintech, 16921-1-AP, 1:100), and mouse anti-RPL4 (Proteintech, 67028-1-Ig, 1:100). Samples were washed with PBS and stained for 1 h at room temperature with secondary antibodies coupled to Alexa Fluor 488 or 568 fluorophores in 10% BSA in PBS buffer.

### In vitro translation assay

Translation competent cell lysate was prepared from HCT116 cells in which the endogenous copies of the eIF3e gene were modified with auxin-inducible degrons allowing rapid depletion of eIF3e by the addition of 500 μM indole-3-acetic acid (IAA) for 12 h ^25^. In vitro translation of nano-luciferase mRNA under the control of the LRRC59 5’UTR with or without the eIF3 binding sites (5’ΔeIF3) was performed with T7 polymerase (NEB). Transcription was performed in the presence of 3′-O-Me-m^7^G(5′)ppp(5′)G RNA Cap Structure Analog (NEB) using linearized plasmid DNA as template and poly-adenylated using polyA polymerase (Invitrogen). RNAs were purified by phenol-chloroform extraction and ethanol precipitation. HCT116 cells were trypsinized and pelleted by centrifugation for 5 min at 1,000 × g at 4 °C, followed by one wash with cold PBS. HCT116 cell pellets were lysed in an equal volume of lysis buffer (10 mM HEPES-KOH pH 7.6, 10 mM KOAc, 0.5 mM Mg(OAc)2, 5 mM DTT, and 1x Complete EDTA-free Proteinase Inhibitor Cocktail tablet (Roche) per 10 ml of buffer). After hypotonic swelling for 45 min on ice, cells were homogenized by forcing it through a syringe attached to a 26G needle approximately 10 times. The lysate was centrifuged at 14,000 × g for 1 min at 4 °C. The supernatant was flash frozen in liquid nitrogen and stored at −80 °C. Each translation reaction contained 50% in vitro translation lysate and buffer to run the final reaction in 0.84 mM ATP, 0.21 mM GTP, 21 mM creatine phosphate (Roche), 45 U ml-1 creatine phosphokinase (Roche), 10 mM HEPES-KOH pH 7.6, 2 mM DTT, 2 mM Mg(OAc)2, 50 mM KOAc, 8 μM amino acids (Promega), 255 μM spermidine, and 1 U/μl murine RNase inhibitor (NEB). Translation reactions were incubated for 1 h at 30 °C. Then translation activity was determined by luciferase assay and confirmed by immunoblotting.

### Fluorescence microscopy

For live-cell imaging and immunofluorescence staining, all samples were visualized by confocal fluorescence microscopy on a Zeiss LSM 980 with Airyscan 2 confocal microscope (Zeiss, Oberkochen, Germany) or a Zeiss LSM5 confocal microscope.

### Fluorescence recovery after photobleaching (FRAP)

For FRAP analysis, MCF7 cells were transfected with the indicated EGFP constructs for 24h. Live cells were viewed under a confocal microscope (Zeiss LSM 980) using a 63× oil immersion objective. After one scanning image, selected regions were then bleached by 3 interactions at 100% laser power (488 diode laser) followed by time-lapse scanning images for 5 min. Mean optical intensity (MOI) of the region of interest was recorded automatically. MOI values within the selected regions were used for calculation. To create fluorescence recovery curves, MOI values were transformed into 0–100% scales of the beginning signal and plotted using Microsoft Excel.

### RT-qPCR

Total RNA from whole cells or RNA retrieved from anti-eIF3b and anti-HSPA8 immunocomplexes was isolated using the Trizol reagent (Life Technology). Quantitative RT-PCR analysis was performed using PerfectStart® Green qPCR SuperMix (TransGen Biotech, cat. AQ601) according to the manufacturer’s instructions. The QuantStudio 3 Real-Time PCR System (ThermoFisher) was used for quantitative analysis. Each target mRNA was quantified in three biological replicates, with each biological replicate having three technical replicates. For primer sequences see Data File S2.

### Crosslinking immunoprecipitation (CLIP) and qPCR

eIF3b-RNA immunocomplexes were prepared following the conditions for the UV-CLIP analysis. Cells were subjected to crosslinking in a UV crosslinker (SCIENTZ, cat. SCIENTZ03-II) set to an energy setting of 400 mJ/cm^2^ for about 4 min. ∼4 × 10^7^ cells were lysed in 1ml lysis buffer (50 mM Tris-HCl, pH 7.4, 100 mM NaCl, 1% NP-40 (Igepal CA630), 0.1% SDS, 0.5% sodium deoxycholate, 1:200 Pierce™ protease inhibitor cocktail (Thermo Scientific, cat. A32955), in RNase/DNase free H_2_O). Lysates were incubated with 30 μl Dynabeads conjugated with 10 μl of anti-eIF3b antibody (Bethyl A301-761A) overnight at 4 °C. After incubation, the supernatant was removed, and beads were washed three times with 1 ml lysis buffer. Bound RNA was isolated by adding Trizol and quantified by qPCR as described above.

### Sucrose density gradient centrifugation

Total lysate was prepared by scraping ∼6 × 10^7^ cells into 0.5 ml hypotonic buffer (5 mM Tris-HCl, pH 7.5, 2.5 mM MgCl^2^, 1.5 mM KCl and 1 x protease inhibitor cocktail) supplemented with 100 μg/ml cycloheximide (CHX), 1 mM DTT, and 100 units of RNAse inhibitor. Triton X-100 and sodium deoxycholate were added to a final concentration of 0.5% each, and samples were vortexed for 5 seconds. Samples were centrifuged at 16,500 g for 7 min at 4°C. Supernatants (cytosolic cell extracts) were collected and absorbance at 260 nm was measured. Approximately 10-15 OD_260_s of lysate were layered over 10% – 50% cold sucrose gradients in buffer (200 mM HEPES-KOH, pH 7.4, 5 mM MgCl_2_, 1 mM KCl, 100 μg/ml CHX, and 1 x protease inhibitor cocktail). Gradients were centrifuged at 39,000 rpm in a Beckman SW28 rotor for 2 h at 4 °C and eluted on a BIOCOMP fractionator system. Fourteen 0.75 ml fractions were collected.

### eIF3-80S crosslinking assay

Polysomal fractions were isolated from MCF7 and AD293 cells seeded to reach 8 × 10^7^ cells on the day of harvest. The cells were treated with 100 μg/ml cycloheximide (CHX) 10 min prior to harvest, then collected into 15 ml Falcon tubes and rinsed once with ice cold PBS. Cells were incubated with 0.5 mM of the crosslinking reagent dithiobis (succinimidyl propionate) (DSP, Sangon Biotech, Cat. #: C110213) and 100 μg/ml CHX in PBS at room temperature for 15 min with rocking. The crosslinking reagent was removed, and cells were incubated with quenching reagent (PBS, 100 μg/ml CHX, 300 mM glycine (Sigma)) for 5 min on ice. Cells were then rinsed again with ice cold PBS and flash frozen in liquid nitrogen.

Cells were lysed in 0.6 ml hypotonic buffer (5 mM Tris–HCl pH 7.5, 2.5 mM MgCl_2_, 1.5 mM KCl, 1 mM DTT, 100 units RNAse inhibitor (Promega, N2111) and 1× Pierce™ protease inhibitor cocktail). Samples were incubated on ice for 30 min and vortexed for 20 s. Triton X-100 and sodium deoxycholate were added to a final concentration of 0.5% each, and lysates were vortexed for another 15 s. Cell lysates were centrifuged at 16,500 g for 7 min at 4 °C. 600 μl supernatant was transferred to an Eppendorf tube and subjected to RNase H digestion by adding the following reagents: 100 units RNase H (NEB), 20 μl of SUPERasIN (ThermoFisher), with a total of 4 μM DNA oligos (IDT), as indicated in the figures. The mixture was incubated at 37 °C for 20 min. After incubation, 10 μl of the RNase H treated lysate mixture was removed to test the efficiency of the RNase H cleavage by RT-qPCR. The remainder of the sample was layered on top of a 10 – 50% cold sucrose gradient in buffer (20 mM HEPES-KOH, pH 7.4, 5 mM MgCl_2_, 100 mM KCl, 100 μg/ml CHX, 10 units/ml RNAse inhibitor, and 1× Pierce™ protease inhibitor cocktail). The gradient was centrifuged at 36,000 rpm (222,000 x g) for 2 h at 4 °C in a SW-41 rotor. After centrifugation, the elution profiles were recorded (BIO-RAD, ECONO UV MONITOR) and polysomal fractions (∼ 5 ml) were collected into a fresh 15 ml Falcon tube using a piston gradient fractionator (BIOCOMP). 100 μl were kept aside for measuring input RNA amounts; the remainder was incubated with 50 μl Dynabeads (Invitrogen) conjugated to 4 μg of anti-eIF3b antibody (Bethyl A301-761A) and incubated with rotation for 16 h at 4 °C. After incubation, beads were washed three times for 10 minutes each with 1 ml CLIP lysis buffer (see above) at room temperature. After the final wash, beads were resuspended in Trizol, RNA was extracted, and RT-qPCR was performed as described above.

### Limited proteolysis

For protease sensitivity assay of LRRC59 and CANX, membrane proteins were extracted from MCF7 cells using ProteoExtract® Membrane protein extraction kit (SIGMA-ALDRICH, Cat. 444810) following the manufacturer’s instructions. For protease sensitivity of TMEM33, the membrane proteins were extracted directly from MCF7 cells using lysis buffer (50 mM Tris-HCl, pH 7.4, 100 mM NaCl, 1% NP-40 (Igepal CA630), 0.1% SDS, 0.5% sodium deoxycholate, 1:200 Pierce™ protease inhibitor cocktail (Thermo Scientific, cat. A32955), in RNase/DNase free H_2_O). 40 μl cell lysate was removed from each sample for zero-minute time points, and digestion was initiated by addition of the amounts of proteinase K indicated in the figures. Reactions were incubated on ice for the durations indicated in the figures. Reactions were stopped by the addition of 1 mM phenylmethylsulphonyl fluoride (PMSF). Samples were analyzed by SDS-PAGE followed by immunoblot with indicated antibodies.

### Quantitative LC-MS/MS

For quantitative proteomics of 80S fractions, MCF7 cell lysate was fractionated on a 10 – 30% sucrose density gradient. 80S monosomes were collected, separated by SDS-PAGE, and stained with Coomassie Blue G-250. The entire gel bands were cut into pieces. Sample were reduced and alkylated with 50 mM ammonium bicarbonate at 37 °C overnight and digested with trypsin. Tryptic peptides were extracted twice with 0.1% formic acid in 50% acetonitrile aqueous solution and dried to reduce volume in a speedvac.

For LC-MS/MS analysis, peptides were separated by a 65 min gradient at a flow rate of 0.300 µl/min with the Thermo EASY-nLC1200 integrated nano-HPLC system which was directly interfaced with the Thermo Exploris 480 mass spectrometer. The analytical column was a home-made fused silica capillary column (75 µm ID, 150 mm length; Upchurch, Oak Harbor, WA) packed with C-18 resin (300 A, 3 µm, Varian, Lexington, MA). Mobile phase A consisted of 0.1% formic acid in water, and mobile phase B consisted of 100% acetonitrile and 0.1% formic acid. The mass spectrometer was operated in the data-dependent acquisition mode using the Xcalibur 4.1 software. A single full-scan mass spectrum was obtained in the Orbitrap (400-1800 m/z, 60,000 resolution), followed by 20 data-dependent MS/MS scans at 30% normalized collision energy. The AGC target was set as 5e4, and the maximum injection time was 50 ms. Each mass spectrum was analyzed using the Thermo Xcalibur Qual Browser and Proteome Discoverer for database searching against the Homo sapiens proteome database downloaded from UniProtKB (UP000005640) containing 80,581 proteins as of October 18, 2022. The Sequest search parameters included a 10 ppm precursor mass tolerance, 0.02 Da fragment ion tolerance, and up to 2 internal cleavage sites. Fixed modifications included cysteine alkylation, and the methionine oxidation was variable modification. Peptides were filtered with 1% false discovery rate (FDR). The mass spectrometry proteomics data have been deposited to the ProteomeXchange Consortium via the PRIDE partner repository with the dataset identifier PXD059488.

## QUANTIFICATION AND STATISTICAL ANALYSIS

P values for the remaining datasets were determined using the multiple t-tests function (two-stage step-up method of Benjamini, Krieger, and Yekutieli) and unpaired t-test in GraphPad Prism 8.

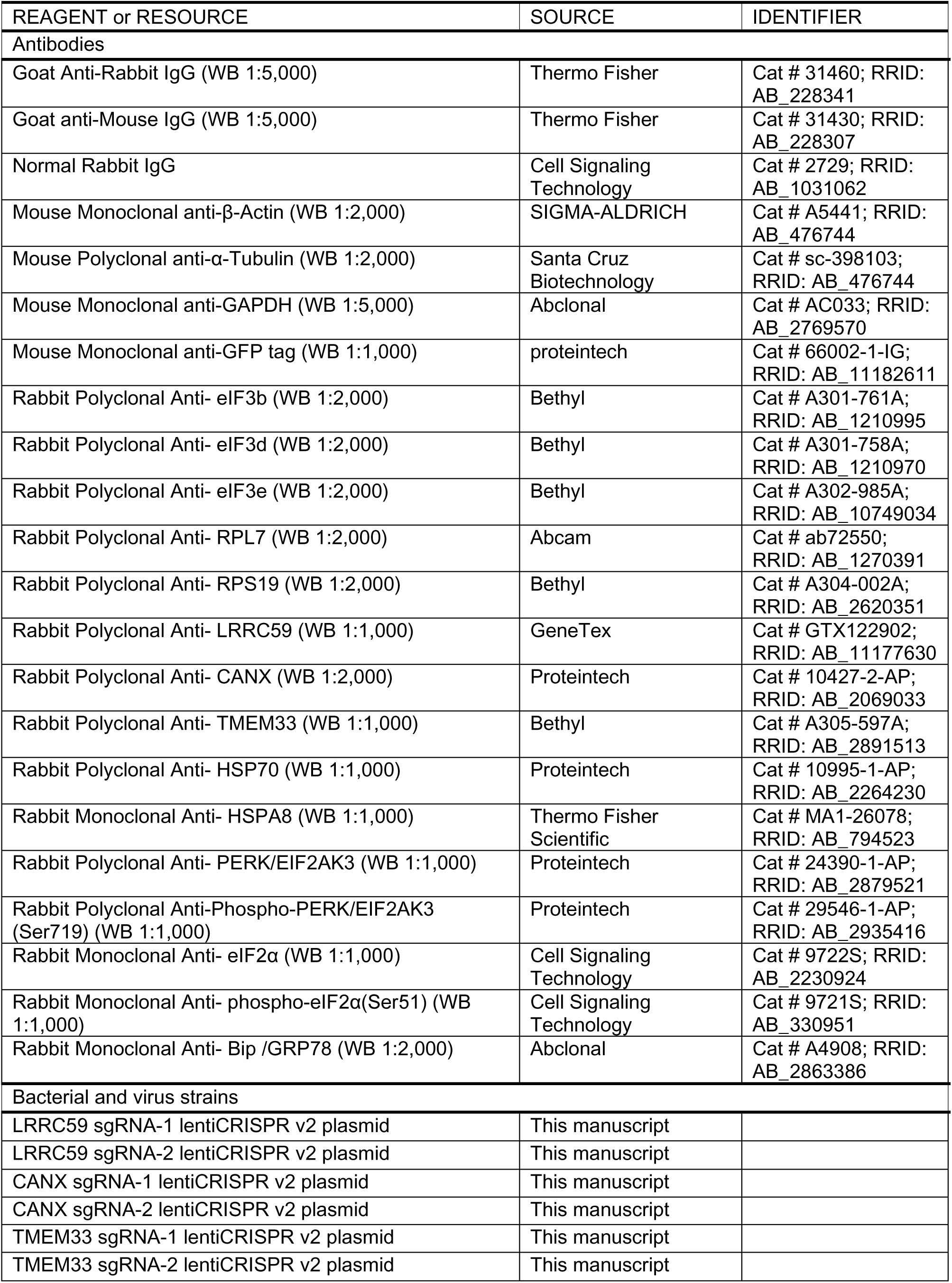
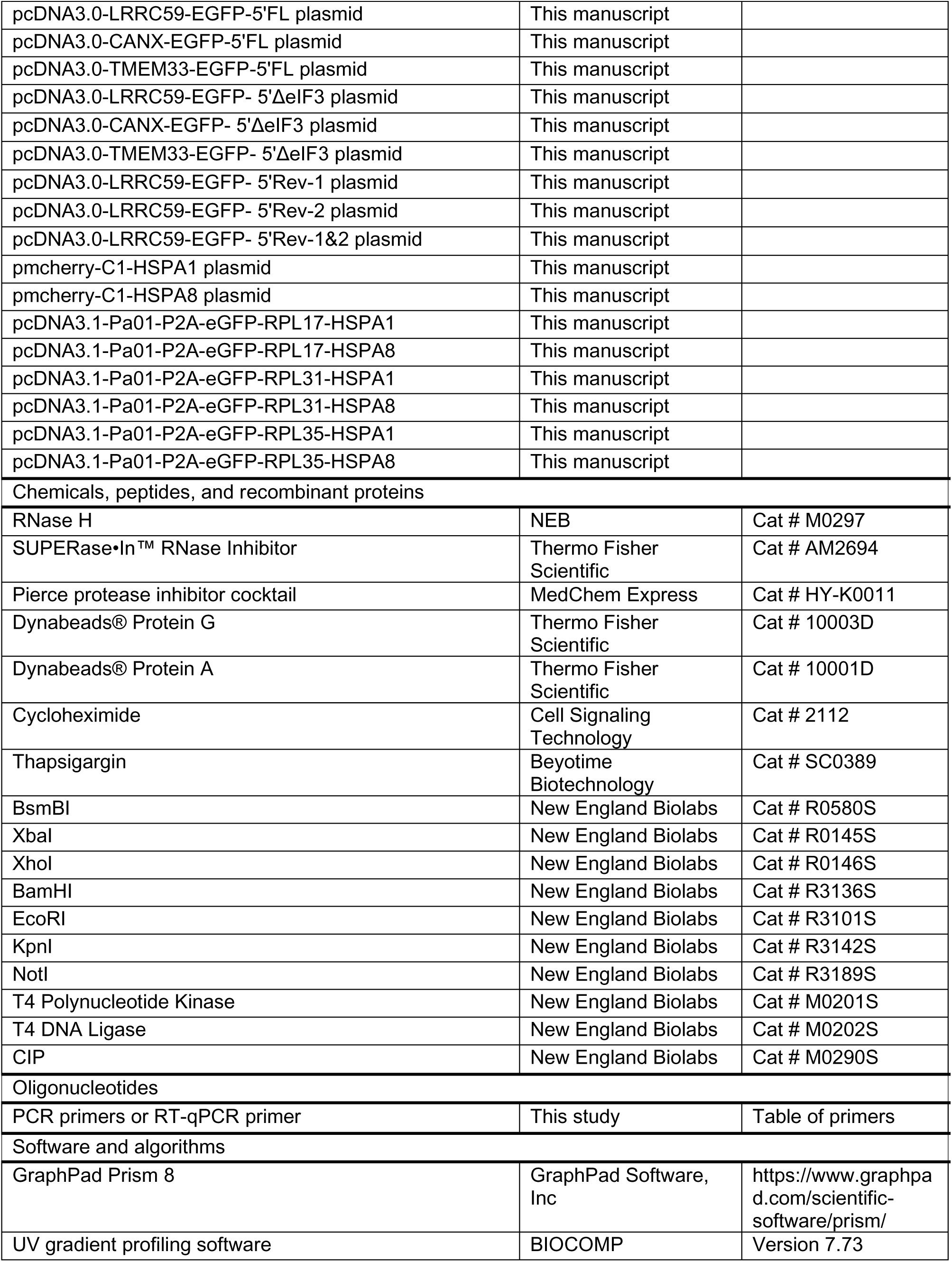
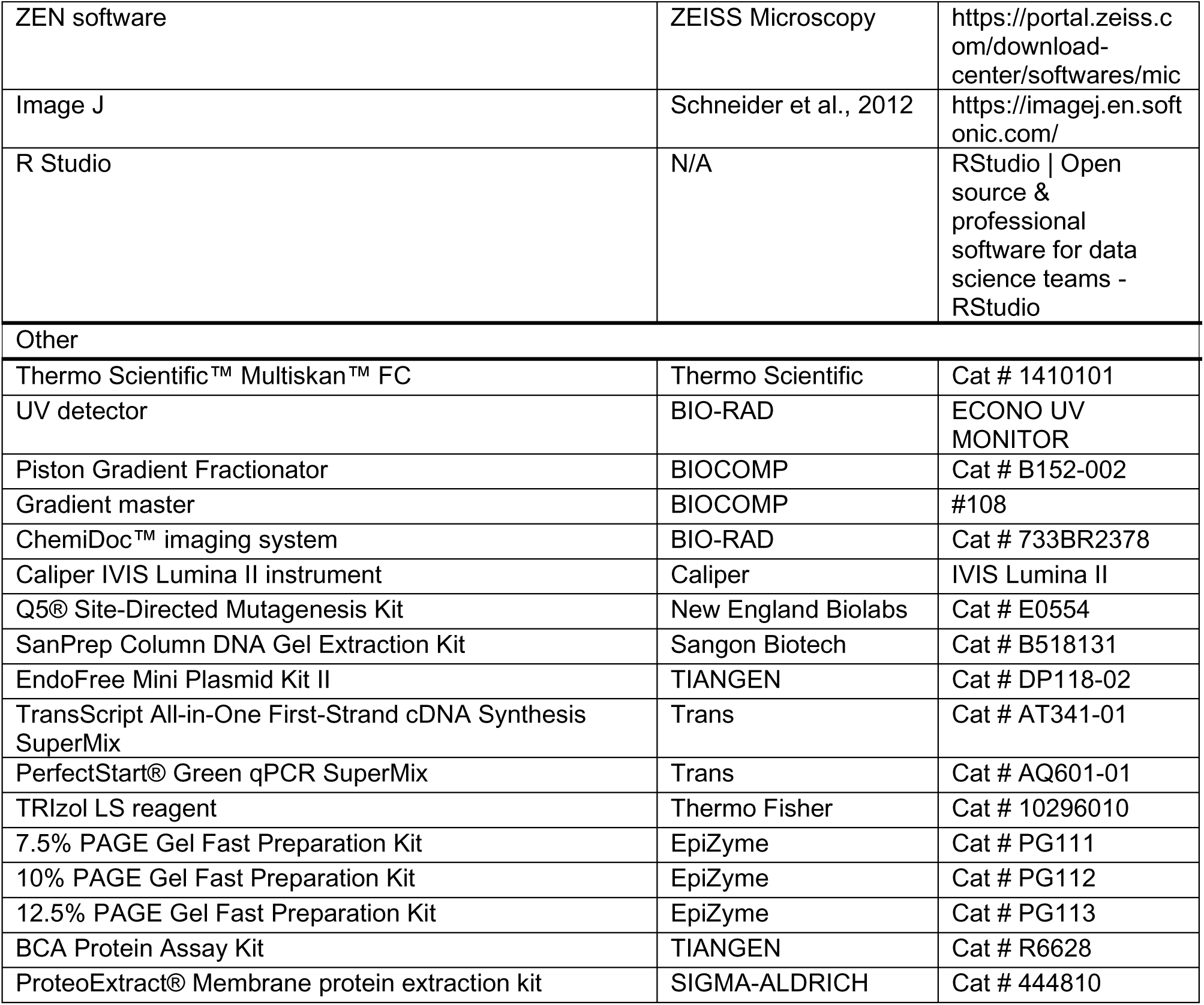

## References

1. Joazeiro, C.A.P. (2019). Mechanisms and functions of ribosome-associated protein quality control. Nature Reviews Molecular Cell Biology 20, 368. 10.1038/s41580-019-0118-2.

2. Kramer, G., Shiber, A., and Bukau, B. (2018). Mechanisms of Cotranslational Maturation of Newly Synthesized Proteins. Annu. Rev. Biochem. 88, 337–364. 10.1146/annurev-biochem-013118-111717.

3. Pechmann, S., Willmund, F., and Frydman, J. (2013). The Ribosome as a Hub for Protein Quality Control. Molecular Cell 49, 411–421. 10.1016/j.molcel.2013.01.020.

4. Wolff, S., Weissman, J.S., and Dillin, A. (2014). Differential Scales of Protein Quality Control. Cell 157, 52–64. 10.1016/j.cell.2014.03.007.

5. Morimoto, R.I. (2008). Proteotoxic stress and inducible chaperone networks in neurodegenerative disease and aging. Genes Dev. 22, 1427–1438. 10.1101/gad.1657108.

6. Thommen, M., Holtkamp, W., and Rodnina, M.V. (2017). Co-translational protein folding: progress and methods. Current Opinion in Structural Biology 42, 83–89. 10.1016/j.sbi.2016.11.020.

7. Döring, K., Ahmed, N., Riemer, T., Suresh, H.G., Vainshtein, Y., Habich, M., Riemer, J., Mayer, M.P., O’Brien, E.P., Kramer, G., et al. (2017). Profiling Ssb-Nascent Chain Interactions Reveals Principles of Hsp70-Assisted Folding. Cell 170, 298–311.e20. 10.1016/j.cell.2017.06.038.

8. Joazeiro, C.A.P. (2019). Mechanisms and functions of ribosome-associated protein quality control. Nat. Rev. Mol. Cell Biol. 20, 368–383. 10.1038/s41580-019-0118-2.

9. Hundley, H.A., Walter, W., Bairstow, S., and Craig, E.A. (2005). Human Mpp11 J Protein: Ribosome-Tethered Molecular Chaperones Are Ubiquitous. Science 308, 1032–1034. 10.1126/science.1109247.

10. Otto, H., Conz, C., Maier, P., Wölfle, T., Suzuki, C.K., Jenö, P., Rücknagel, P., Stahl, J., and Rospert, S. (2005). The chaperones MPP11 and Hsp70L1 form the mammalian ribosome-associated complex. PNAS 102, 10064–10069. 10.1073/pnas.0504400102.

11. Jaiswal, H., Conz, C., Otto, H., Wolfle, T., Fitzke, E., Mayer, M.P., and Rospert, S. (2011). The Chaperone Network Connected to Human Ribosome-Associated Complex. Molecular and Cellular Biology 31, 1160–1173. 10.1128/MCB.00986-10.

12. Tian, G., Hu, C., Yun, Y., Yang, W., Dubiel, W., Cheng, Y., and Wolf, D.A. (2021). Dual roles of HSP70 chaperone HSPA1 in quality control of nascent and newly synthesized proteins. EMBO J 40, e106183. 10.15252/embj.2020106183.

13. Willmund, F., del Alamo, M., Pechmann, S., Chen, T., Albanèse, V., Dammer, E.B., Peng, J., and Frydman, J. (2013). The Cotranslational Function of Ribosome-Associated Hsp70 in Eukaryotic Protein Homeostasis. Cell 152, 196–209. 10.1016/j.cell.2012.12.001.

14. Stein, K.C., Kriel, A., and Frydman, J. (2019). Nascent Polypeptide Domain Topology and Elongation Rate Direct the Cotranslational Hierarchy of Hsp70 and TRiC/CCT. Molecular Cell 75, 1117–1130.e5. 10.1016/j.molcel.2019.06.036.

15. Gumiero, A., Conz, C., Gesé, G.V., Zhang, Y., Weyer, F.A., Lapouge, K., Kappes, J., von Plehwe, U., Schermann, G., Fitzke, E., et al. (2016). Interaction of the cotranslational Hsp70 Ssb with ribosomal proteins and rRNA depends on its lid domain. Nat Commun 7, 13563. 10.1038/ncomms13563.

16. Lee, K., Ziegelhoffer, T., Delewski, W., Berger, S.E., Sabat, G., and Craig, E.A. (2021). Pathway of Hsp70 interactions at the ribosome. Nat Commun 12, 5666. 10.1038/s41467-021-25930-8.

17. Lin, Y., Li, F., Huang, L., Polte, C., Duan, H., Fang, J., Sun, L., Xing, X., Tian, G., Cheng, Y., et al. (2020). eIF3 Associates with 80S Ribosomes to Promote Translation Elongation, Mitochondrial Homeostasis, and Muscle Health. Molecular Cell 79, 575–587.e7. 10.1016/j.molcel.2020.06.003.

18. Sha, Z., Brill, L.M., Cabrera, R., Kleifeld, O., Scheliga, J.S., Glickman, M.H., Chang, E.C., and Wolf, D.A. (2009). The eIF3 Interactome Reveals the Translasome, a Supercomplex Linking Protein Synthesis and Degradation Machineries. Molecular Cell 36, 141–152. 10.1016/j.molcel.2009.09.026.

19. Hinnebusch, A.G. (2017). Structural Insights into the Mechanism of Scanning and Start Codon Recognition in Eukaryotic Translation Initiation. Trends in Biochemical Sciences 42, 589–611. 10.1016/j.tibs.2017.03.004.

20. Valášek, L.S., Zeman, J., Wagner, S., Beznosková, P., Pavlíková, Z., Mohammad, M.P., Hronová, V., Herrmannová, A., Hashem, Y., and Gunišová, S. (2017). Embraced by eIF3: structural and functional insights into the roles of eIF3 across the translation cycle. Nucleic Acids Res 45, 10948–10968. 10.1093/nar/gkx805.

21. Wolf, D.A., Lin, Y., Duan, H., and Cheng, Y. (2020). eIF-Three to Tango: emerging functions of translation initiation factor eIF3 in protein synthesis and disease. J Mol Cell Biol 12, 403–409. 10.1093/jmcb/mjaa018.

22. Cate, J.H.D. (2017). Human eIF3: from ‘blobology’ to biological insight. Phil. Trans. R. Soc. B 372, 20160176. 10.1098/rstb.2016.0176.

23. Wagner, S., Herrmannová, A., Malík, R., Peclinovská, L., and Valášek, L.S. (2014). Functional and Biochemical Characterization of Human Eukaryotic Translation Initiation Factor 3 in Living Cells. Mol. Cell. Biol. 34, 3041–3052. 10.1128/MCB.00663-14.

24. Wagner, S., Herrmannová, A., Šikrová, D., and Valášek, L.S. (2016). Human eIF3b and eIF3a serve as the nucleation core for the assembly of eIF3 into two interconnected modules: the yeast-like core and the octamer. Nucl. Acids Res., gkw972. 10.1093/nar/gkw972.

25. Duan, H., Zhang, S., Zarai, Y., Öllinger, R., Wu, Y., Sun, L., Hu, C., He, Y., Tian, G., Rad, R., et al. (2023). eIF3 mRNA selectivity profiling reveals eIF3k as a cancer-relevant regulator of ribosome content. The EMBO Journal, e112362. 10.15252/embj.2022112362.

26. Peterson, D.T., Merrick, W.C., and Safer, B. (1979). Binding and release of radiolabeled eukaryotic initiation factors 2 and 3 during 80 S initiation complex formation. J. Biol. Chem. 254, 2509–2516.

27. Trachsel, H., and Staehelin, T. (1979). Initiation of mammalian protein synthesis The multiple functions of the initiation factor eIF-3. Biochimica et Biophysica Acta (BBA) - Nucleic Acids and Protein Synthesis 565, 305–314. 10.1016/0005-2787(79)90207-7.

28. Bohlen, J., Fenzl, K., Kramer, G., Bukau, B., and Teleman, A.A. (2020). Selective 40S Footprinting Reveals Cap-Tethered Ribosome Scanning in Human Cells. Molecular Cell 79, 561–574.e5. 10.1016/j.molcel.2020.06.005.

29. Wagner, S., Herrmannová, A., Hronová, V., Gunišová, S., Sen, N.D., Hannan, R.D., Hinnebusch, A.G., Shirokikh, N.E., Preiss, T., and Valášek, L.S. (2020). Selective Translation Complex Profiling Reveals Staged Initiation and Co-translational Assembly of Initiation Factor Complexes. Molecular Cell 79, 546–560.e7. 10.1016/j.molcel.2020.06.004.

30. Shu, X.E., Mao, Y., Jia, L., and Qian, S.-B. (2021). Dynamic eIF3a O-GlcNAcylation controls translation reinitiation during nutrient stress. Nat Chem Biol, 1–8. 10.1038/s41589-021-00913-4.

31. Lee, A.S.Y., Kranzusch, P.J., and Cate, J.H.D. (2015). eIF3 targets cell-proliferation messenger RNAs for translational activation or repression. Nature 522, 111–114. 10.1038/nature14267.

32. Meyer, K.D., Patil, D.P., Zhou, J., Zinoviev, A., Skabkin, M.A., Elemento, O., Pestova, T.V., Qian, S.-B., and Jaffrey, S.R. (2015). 5′ UTR m6A Promotes Cap-Independent Translation. Cell 163, 999–1010. 10.1016/j.cell.2015.10.012.

33. Lu, Z., Zhang, Q.C., Lee, B., Flynn, R.A., Smith, M.A., Robinson, J.T., Davidovich, C., Gooding, A.R., Goodrich, K.J., Mattick, J.S., et al. (2016). RNA Duplex Map in Living Cells Reveals Higher-Order Transcriptome Structure. Cell 165, 1267–1279. 10.1016/j.cell.2016.04.028.

34. Zubradt, M., Gupta, P., Persad, S., Lambowitz, A.M., Weissman, J.S., and Rouskin, S. (2017). DMS-MaPseq for genome-wide or targeted RNA structure probing in vivo. Nat Methods 14, 75–82. 10.1038/nmeth.4057.

35. Chen, C.R., Li, Y.C., Chen, J., Hou, M.C., Papadaki, P., and Chang, E.C. (1999). Moe1, a conserved protein in Schizosaccharomyces pombe, interacts with a Ras effector, Scd1, to affect proper spindle formation. Proc. Natl. Acad. Sci. U.S.A. 96, 517–522.

36. Shah, M., Su, D., Scheliga, J.S., Pluskal, T., Boronat, S., Motamedchaboki, K., Campos, A.R., Qi, F., Hidalgo, E., Yanagida, M., et al. (2016). A Transcript-Specific eIF3 Complex Mediates Global Translational Control of Energy Metabolism. Cell Reports 16, 1891–1902. 10.1016/j.celrep.2016.07.006.

37. Dickson, K.M., Bergeron, J.J.M., Shames, I., Colby, J., Nguyen, D.T., Chevet, E., Thomas, D.Y., and Snipes, G.J. (2002). Association of calnexin with mutant peripheral myelin protein-22 ex vivo: A basis for “gain-of-function” ER diseases. PNAS 99, 9852–9857. 10.1073/pnas.152621799.

38. Hewett, J., Gonzalez-Agosti, C., Slater, D., Ziefer, P., Li, S., Bergeron, D., Jacoby, D.J., Ozelius, L.J., Ramesh, V., and Breakefield, X.O. (2000). Mutant torsinA, responsible for early-onset torsion dystonia, forms membrane inclusions in cultured neural cells. Human Molecular Genetics 9, 1403–1413. 10.1093/hmg/9.9.1403.

39. Xu, F., Du, W., Zou, Q., Wang, Y., Zhang, X., Xing, X., Li, Y., Zhang, D., Wang, H., Zhang, W., et al. (2021). COPII mitigates ER stress by promoting formation of ER whorls. Cell Res 31, 141–156. 10.1038/s41422-020-00416-2.

40. Guo, Y., Shen, D., Zhou, Y., Yang, Y., Liang, J., Zhou, Y., Li, N., Liu, Y., Yang, G., and Li, W. (2022). Deep Learning-Based Morphological Classification of Endoplasmic Reticulum Under Stress. Frontiers in Cell and Developmental Biology 9.

41. Albanèse, V., Yam, A.Y.-W., Baughman, J., Parnot, C., and Frydman, J. (2006). Systems Analyses Reveal Two Chaperone Networks with Distinct Functions in Eukaryotic Cells. Cell 124, 75–88. 10.1016/j.cell.2005.11.039.

42. McCallum, C.D., Do, H., Johnson, A.E., and Frydman, J. (2000). The Interaction of the Chaperonin Tailless Complex Polypeptide 1 (Tcp1) Ring Complex (Tric) with Ribosome-Bound Nascent Chains Examined Using Photo-Cross-Linking. The Journal of Cell Biology 149, 591–602. 10.1083/jcb.149.3.591.

43. Thulasiraman, V. (1999). In vivo newly translated polypeptides are sequestered in a protected folding environment. The EMBO Journal 18, 85–95. 10.1093/emboj/18.1.85.

44. Schlecht, R., Scholz, S.R., Dahmen, H., Wegener, A., Sirrenberg, C., Musil, D., Bomke, J., Eggenweiler, H.-M., Mayer, M.P., and Bukau, B. (2013). Functional Analysis of Hsp70 Inhibitors. PLoS ONE 8, e78443. 10.1371/journal.pone.0078443.

45. Kišonaitė, M., Wild, K., Lapouge, K., Gesé, G.V., Kellner, N., Hurt, E., and Sinning, I. (2023). Structural inventory of cotranslational protein folding by the eukaryotic RAC complex. Nat Struct Mol Biol 30, 670–677. 10.1038/s41594-023-00973-1.

46. Chen, Y., Tsai, B., Li, N., and Gao, N. (2022). Structural remodeling of ribosome associated Hsp40-Hsp70 chaperones during co-translational folding. Nat Commun 13, 3410. 10.1038/s41467-022-31127-4.

47. De Silva, D., Ferguson, L., Chin, G.H., Smith, B.E., Apathy, R.A., Roth, T.L., Blaeschke, F., Kudla, M., Marson, A., Ingolia, N.T., et al. (2021). Robust T cell activation requires an eIF3-driven burst in T cell receptor translation. eLife 10, e74272. 10.7554/eLife.74272.

48. Petrychenko, V., Yi, S.-H., Liedtke, D., Peng, B.-Z., Rodnina, M.V., and Fischer, N. (2024). Structural basis for translational control by the human 48S initiation complex. Nat Struct Mol Biol, 1–11. 10.1038/s41594-024-01378-4.

49. Chartron, J.W., Hunt, K.C.L., and Frydman, J. (2016). Cotranslational signal-independent SRP preloading during membrane targeting. Nature 536, 224–228. 10.1038/nature19309.

50. Ma, W., and Mayr, C. (2018). A Membraneless Organelle Associated with the Endoplasmic Reticulum Enables 3′UTR-Mediated Protein-Protein Interactions. Cell 175, 1492–1506.e19. 10.1016/j.cell.2018.10.007.

51. Pulos-Holmes, M.C., Srole, D.N., Juarez, M.G., Lee, A.S.-Y., McSwiggen, D.T., Ingolia, N.T., and Cate, J.H. (2019). Repression of ferritin light chain translation by human eIF3. eLife 8, e48193. 10.7554/eLife.48193.

52. Lee, A.S.Y., Kranzusch, P.J., Doudna, J.A., and Cate, J.H.D. (2016). eIF3d is an mRNA cap-binding protein that is required for specialized translation initiation. Nature 536, 96–99. 10.1038/nature18954.

53. Mukhopadhyay, S., Amodeo, M.E., and Lee, A.S.Y. (2023). eIF3d controls the persistent integrated stress response. Molecular Cell, S1097276523006433. 10.1016/j.molcel.2023.08.008.

54. Walker, M.J., Shortridge, M.D., Albin, D.D., Cominsky, L.Y., and Varani, G. (2020). Structure of the RNA Specialized Translation Initiation Element that Recruits eIF3 to the 5′-UTR of c-Jun. Journal of Molecular Biology 432, 1841–1855. 10.1016/j.jmb.2020.01.001.

55. Natsume, T., Kiyomitsu, T., Saga, Y., and Kanemaki, M.T. (2016). Rapid Protein Depletion in Human Cells by Auxin-Inducible Degron Tagging with Short Homology Donors. Cell Reports 15, 210–218. 10.1016/j.celrep.2016.03.001.

56. Zhao, W., Zhang, S., Zhu, Y., Xi, X., Bao, P., Ma, Z., Kapral, T.H., Chen, S., Zagrovic, B., Yang, Y.T., et al. (2022). POSTAR3: an updated platform for exploring post-transcriptional regulation coordinated by RNA-binding proteins. Nucleic Acids Research 50, D287–D294. 10.1093/nar/gkab702.

