## Supplemental Information for "5’UTR-mediated retention of eIF3 on 80S ribosomes promotes co-translational folding of ER membrane proteins"

- Figures S1 – S6
- Supplementary Data File S1, S2
- Supplementary Movie 1 - 3

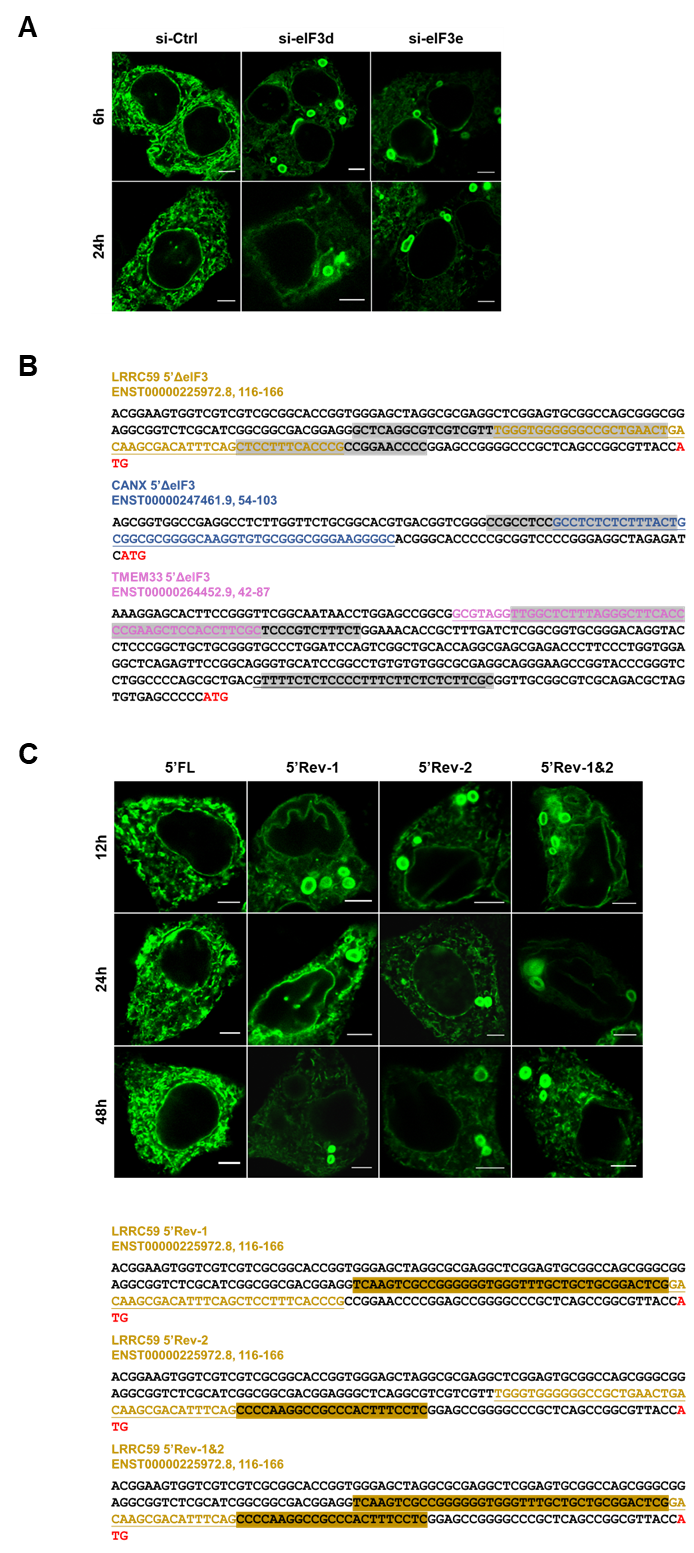

**Supplementary Figure 1 Effect of eIF3 on the localization of LRRC59** (related to Figure 1)

**A.** LRRC59-EGFP under control of its cognate 5’UTR (Figure 1A) was expressed in MCF7 cells treated with control siRNA (si-Ctrl) or siRNA targeting eIF3d or eIF3e (si-eIF3e) for 72 h. 6 and 24 h after transfection, EGFP fusion proteins were detected by fluorescence microscopy of live cells. Scale bars: 5 μm.

**B.** Sequences of the 5’UTRs of mRNAs encoding the indicated membrane proteins. The underlined sequences represent the eIF3 binding sites mapped by CLIP. The grey areas highlight the sequences deleted from plasmids expressing the membrane proteins as fusions with EGFP. Start codons are highlighted in red font. The 5’UTR of *TMEM33* contains a second potential eIF3 binding site (underlined) which was considered minor based on the rarity of corresponding CLIP fragments. The site was deleted in the expression plasmid as shown but was not edited by CRISPR as shown in Figure 2A.

**C.** The eIF3 binding sites in plasmids driving the expression of LRRC59 fusions with EGFP under control of the cognate 5’UTR were altered by inverting the half sites of the stem loop shown in Figure 1C either individually (5’Rev-1, 5’Rev-2) or simultaneously (5’Rev-1&2). The plasmids were transfected into MCF7 cells, and live cells were imaged after 12 – 24 h. Scale bars: 5 μm.

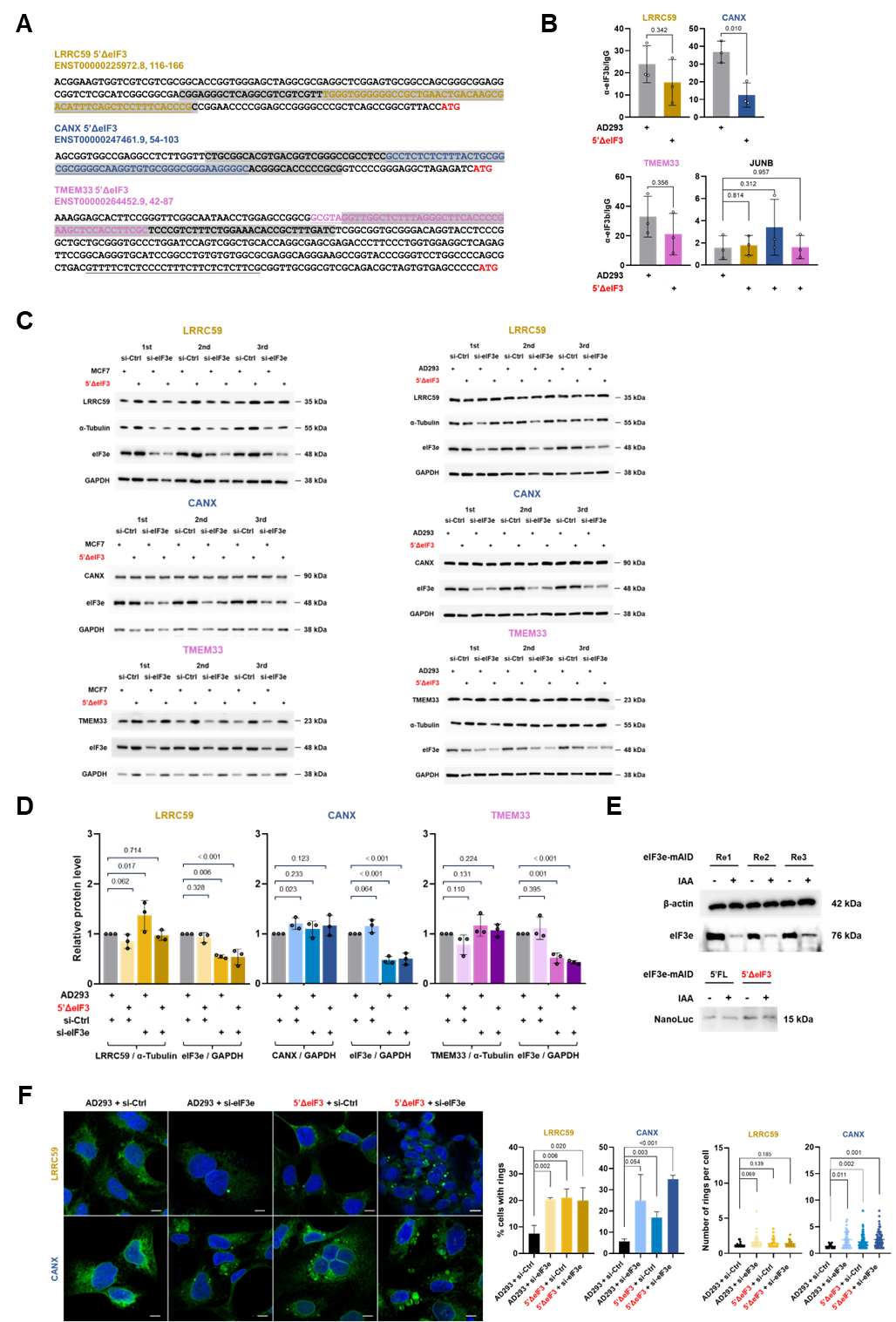

**Supplementary Figure 2** **Effect of eIF3 5’UTR binding sites on the localization of ER membrane proteins in AD293 cells** (related to Figure 2)

**A.** Sequences of the 5’UTRs of mRNAs encoding the indicated membrane proteins. The underlined sequences represent the eIF3 binding sites mapped by CLIP. The grey areas highlight the sequences deleted by genome editing in AD293 cells. Start codons are highlighted in red font. The 5’UTR of *TMEM33* contains a second potential eIF3 binding site (underlined) which was considered minor based on the rarity of corresponding CLIP fragments and was therefore not edited.

**B.** mRNPs present in cell lysate from parental AD293 cells or from AD293 cells in which the indicated eIF3 binding regions were deleted (5’ΔeIF3 cell lines) were immunopurified with eIF3b antibodies. Co-purification of the indicated mRNAs was quantified by RT-qPCR. Bars represent means (n = 3) ± standard deviations, numbers represent p values (two-stage step-up method of Benjamini, Krieger, and Yekutieli).

**C.** Cell lysates prepared from MCF7 (left) and AD293 (right) cells deleted of eIF3 5’UTR binding sites (5’ΔeIF3) within genes encoding the ER membrane proteins LRRC59, CANX, and TMEM33 were analyzed by immunoblotting with the specified antibodies. Where indicated, cells were transfected with nontargeting control siRNA (si-Ctrl) or with siRNA knocking down eIF3e for 72 h. Triplicate datasets are shown.

**D.** The Western blots in C. were quantified. The signals of ER membrane proteins were normalized to reference proteins (GAPDH or α-tubulin) as indicated. Bars represent means (n = 3) ± standard deviations, numbers represent p values (two-stage step-up method of Benjamini, Krieger, and Yekutieli).

**E.** In vitro translation of nano-luciferase mRNA under the control of the *LRRC59* 5’UTR with or without the eIF3 binding sites (5’ΔeIF3). Translation competent cell lysate was prepared from HCT116 cells in which the endogenous copies of the eIF3e gene were modified with auxin-inducible degrons allowing rapid depletion of eIF3e by the addition of 500 μM indole-3-acetic acid (IAA) for 12 h ^1^. The efficiency of eIF3e depletion and translation activity were determined by immunoblotting.

**F.** Parental AD293 cells or AD293 cells in which the indicated eIF3 binding regions were deleted (5’ΔeIF3 cell lines) and/or eIF3e was knocked down for 72 h were fixed and stained with antibodies against LRRC59 or CANX. Bright ring-like structures were visualized by confocal microscopy, and the percentage of ring containing cells were scored (top panels). Bars represent means (n = >200 cells) ± standard deviations, numbers represent p values (two-stage step-up method of Benjamini, Krieger, and Yekutieli). In addition, the number of rings per cell was scored (bottom panels) with data representing means (n = >39 cells) ± standard deviations and numbers representing p values (unpaired t-test). Scale bars: 10 μm.

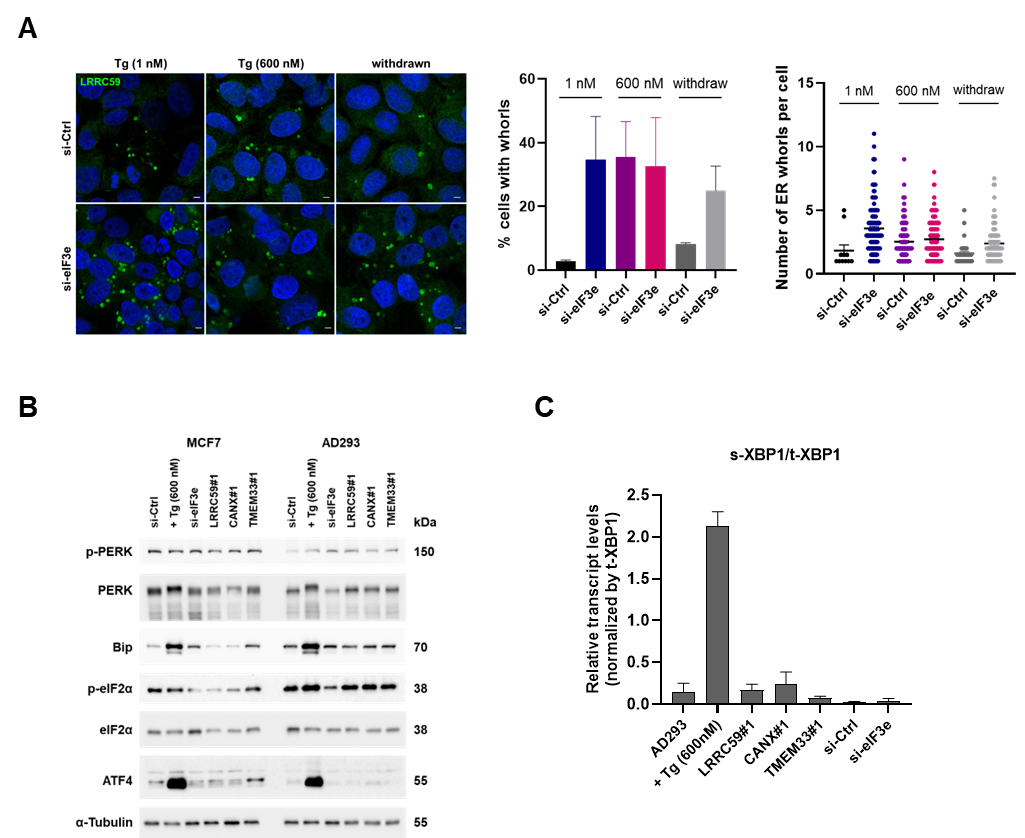

**Supplementary Figure 3 Effect of disabling eIF3 on ER stress markers** (related to Figure 3)

**A.** MCF7 cells transfected with control siRNA (si-Ctrl) or siRNA targeting eIF3e (si-eIF3e) for 72 h were treated with the indicated concentrations of the ER stress inducer thapsigargin (Tg) for 6 h, followed by withdrawal of Tg for 6 hours. LRRC59 positive ER whorls were detected by indirect immunofluorescence staining and visualized by confocal microscopy, and the percentage of ring containing cells were scored (top panels). Bars represent means (n = >200 cells) ± standard deviations, numbers represent p values (two-stage step-up method of Benjamini, Krieger, and Yekutieli). In addition, the number of rings per cell was scored (bottom panels) with data representing means (n = >39 cells) ± standard deviations and numbers representing p values (unpaired t-test). Scale bars: 10 μm.

**B.** Parental MCF7 and AD293 cells were treated with thapsigargin for 6 h or transfected with control siRNA (si-Ctrl) or siRNA targeting eIF3e (si-eIF3e) for 72 h, followed by preparation of cell lysates. In parallel, lysates were prepared from MCF7 and AD293 cells deleted of eIF3 5’UTR binding sites (5’ΔeIF3) within genes encoding the ER membrane proteins LRRC59, CANX, and TMEM33. Cell lysates were subjected to immunoblotting with antibodies against ER stress markers. Alpha-tubulin levels are shown for reference.

**C.** RNA was prepared from MCF7 cells treated as in (B.) and analyzed by RT-qPCR for the level of splicing of *XBP1* mRNA (*s-XPB1*). The graph shows *s-XBP1* mRNA levels relative to total *XBP1* mRNA (t-*XBP1*). Bars represent averages (n = 3) ± standard deviations. Numbers represent p values (two-stage step-up method of Benjamini, Krieger, and Yekutieli).

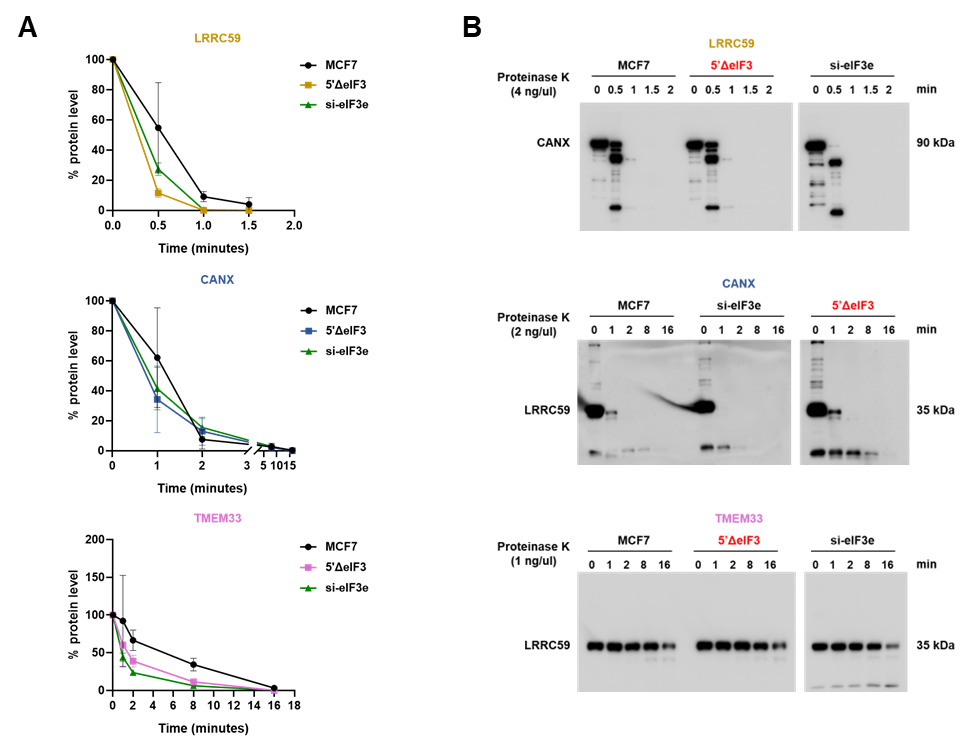

**Supplementary Figure 4** **Effect of eIF3 deficiency on the folding state of ER membrane proteins** (related to Figure 4)

**A.** Quantification of the blots in Figure 4D. – F. showing the sensitivity of ER membrane proteins to limited proteolysis in lysate from parental MCF7 cells or from MCF7 cells in which the indicated eIF3 binding regions were deleted (5’ΔeIF3 cell lines) or eIF3e was knocked down for 72 h. In each case, the full-length proteins present at t = 0 were quantified. The line graphs show averages (n = 3) ± standard deviations.

**B.** The protein lysates used in Figure 4D. – F. showing the sensitivity of ER membrane proteins to limited proteolysis depending on eIF3 function were probed with antibodies against ER membrane proteins whose encoding mRNAs were not edited in the respective cell lines. On the top panel, the protease sensitivity of CANX was assessed in cells in which the eIF3 binding site was deleted from the 5’UTR of *LRRC59* mRNA (LRRC59 5’ΔeIF3). As expected, whereas LRRC59 is hypersensitive to proteinase K in both eIF3e knockdown and 5’ΔeIF3 cells, CANX is only hypersensitive in eIF3e knockdown cells. The middle and the bottom panels show similar specificity controls for CANX and TMEM33 5’ΔeIF3 cell lines.

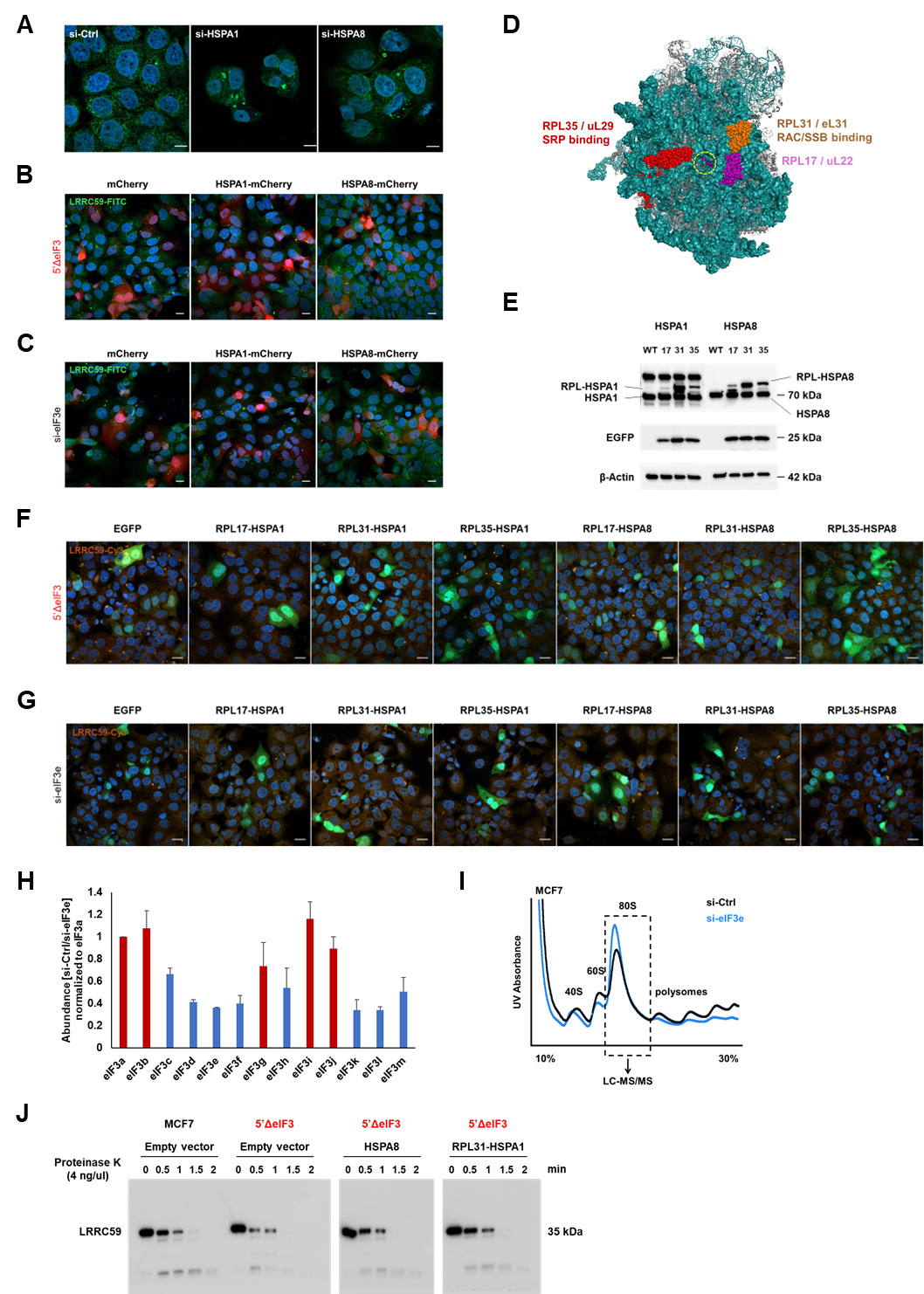

**Supplementary Figure 5** **Effect of eIF3 on chaperone-mediated folding of LRRC59** (related to Figure 5)

**A.** LRRC59 positive ER whorls in MCF7 cells transfected with control siRNA (si-Ctrl) or siRNA targeting HSPA1 or HSPA8 for 72 h. Due to severe cytotoxicity of HSP70 knockdown in MCF7 cells, too few cells were available for numerical scoring and statistical analysis.

**B., C.** MCF7 cells in which the eIF3 binding region was deleted from the 5’UTR of *LRRC59* (B.) or that were transfected with si-eIF3e for 72 h (C.) were transfected with plasmids driving the expression of mCherry or mCherry fused to the C-termini of HSPA1 or HSPA8 for 48 h. Cells were fixed, and LRRC59 was detected by immunofluorescence staining. Samples were imaged by confocal microscopy and representative overlays of LRRC59 (green), mCherry fusions (red), and DAPI stained cell nuclei (blue) are shown. Scale bars 20 μm. The percentage of cells showing LRRC59 positive ER whorls were scored (Figure 5C., D.)

**D.** Position of 60S ribosomal proteins according to PDB entry 4UG0 ^2^. The ribosome exit tunnel is indicated by the stippled circle.

**E.** The 60S ribosomal proteins highlighted in D. were fused to the N-termini of HSPA1 and HSPA8. Fusion proteins were expressed in MCF7 cells in which the eIF3 binding regions in the 5’UTRs of LRRC59 were deleted (5’ΔeIF3 cell lines) or eIF3e was knocked down for 72 h. Cells also express EGFP which was separated from RPL fusions by a P2A protease cleavage site. Cell lysates were subjected to immunoblotting to detect the indicated RPL fusion proteins and EGFP.

**F., G.** Same experiment as in E. but cells were fixed and LRRC59 positive ER whorls were detected by indirect immunofluorescence. Samples were imaged by confocal microscopy and representative overlays of LRRC59 (red), EGFP (green), and DAPI stained cell nuclei (blue) are shown. Scale bars 20 μm. LRRC59 positive ER whorls were quantitatively scored (Figure 5E., F.).

**H.** Cell lysates from MCF7 cells transfected with control (si-Ctrl) or eIF3e targeting siRNA (si-eIF3e) were separated by sucrose density gradient centrifugation (see I.). Fractions containing 80S ribosomes were analyzed by quantitative LC-MS/MS. The relative abundance of eIF3 subunits in 80S fractions of Ctrl versus eIF3e knockdown cells was plotted in a bar graph (averages of n = 2).

**I.** UV595 traces of the sucrose density gradients. The boxed region shows the 80S fraction that was analyzed by quantitative LC-MS/MS.

**J.** Parental MCF7 cells or MCF7 cells in which the eIF3 binding region in the LRRC59 gene were deleted (5’ΔeIF3 cell line) were transfected with HSPA8-mCherry, RPL31-HSPA1, or empty vector control. LRRC59 was extracted and subjected to limited proteolysis with proteinase K, followed by detection of proteolytic products by immunoblotting.

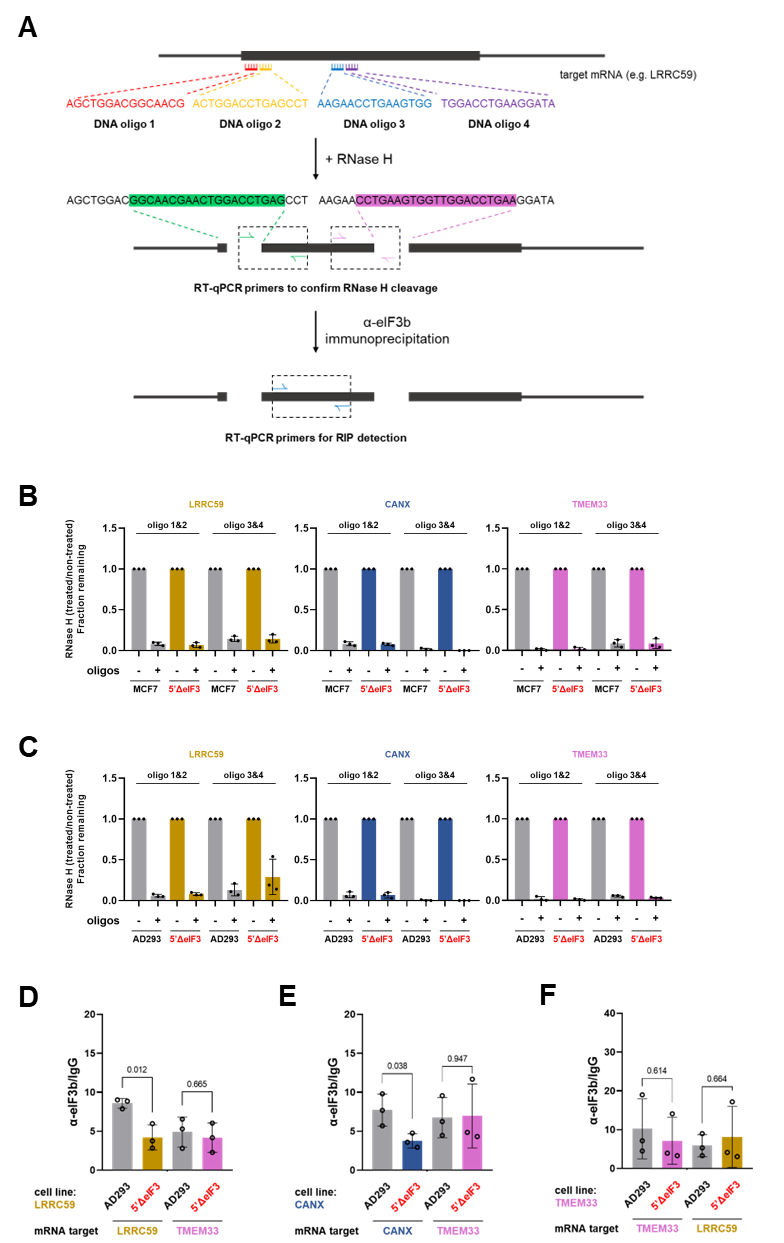

**Supplementary Figure 6** **Effect of 5’UTR eIF3 binding sites on the stability of start codon proximal eIF3-80S** (related to Figure 6)

**A.** Diagram of oligonucleotide-mediated cleavage of start codon proximal regions of mRNAs encoding ER membrane proteins. Following DSP cross-linking, RNAse H was targeted by two sets of adjacent oligonucleotides at either end of the targeted mRNA regions (upstream oligos 1&2, downstream oligos 3&4). Cleavage efficiency was determined separately for the upstream and downstream cleavage sites by RT-qPCR using two sets of PCR primers (upstream PCR #1, downstream PCR #2).

**B.** Determination of RNAse H cleavage efficiency. The above described RNAse H cleavage and RT-qPCR reactions were carried out with RNA samples isolated from parental MCF7 cells and MCF7 cells in which the eIF3 5’UTR binding sites of LRRC59, CANX, or TMEM33 were deleted by CRISPR-mediated gene editing (5’ΔeIF3). Bars represent fractions remaining after RNAse H cleavage (n = 3) ± standard deviations.

**C.** Same as in B. but with parental and 5’ΔeIF3 AD293 cells.

**D.** Cross-linking assay to measure the abundance of eIF3-80S complexes between codons 24 - 107 of *LRRC59* mRNA. Lysate from parental AD293 cells and AD293 cells in which the eIF3 binding region was deleted from the 5’UTR of *LRRC59* were used for anti-eIF3b or IgG (negative control) IP. The retained mRNA fragments were detected by RT-qPCR with primers specific for *LRRC59* (2 bars on the left) or *TMEM33* mRNA (specificity control, 2 bars on the right). Bars represent average enrichment of mRNA in anti-eIF3b IPs relative to IgG (n = 3, ± standard deviations); numbers represent p values (two-stage step-up method of Benjamini, Krieger, and Yekutieli).

**E.** Cross-linking assay to measure the abundance of eIF3-80S complexes between codons 19 - 119 of *CANX* mRNA. Same experiment as in D. but with AD293 cells in which the eIF3 binding region was deleted from the 5’UTR of *CANX.* *TMEM33* was used as a specificity control. Bar graph as in D.

**F.** Cross-linking assay to measure the abundance of eIF3-80S complexes between codons 32 – 76 of *TMEM33* mRNA. Same experiment as in D. but with AD293 cells in which the eIF3 binding region was deleted from the 5’UTR of *TMEM33.* *LRRC59* was used as a specificity control. Bar graph as in D.

**Supplementary Data File S1** (related to Figure 5)

Proteomic data shown in Figure 5G.

**Supplementary Data File S2**

Sequences of all oligonucleotides used in this study

**Supplementary Movie 1. FRAP assay with LRRC59-EGFP** (Related to Figure 4)

MCF7 cells were transfected with plasmids encoding the LRRC59-EGFP fusion proteins under control of its cognate 5’UTRs from which the eIF3 binding site was deleted (5’ΔeIF3, **Figure S1B**). After 24 h, live cells were photobleached in the indicated areas of tubular ER or ER whorls, and fluorescence recovery was observed over time. Scale bar: 5 μm.

**Supplementary Movie 2. FRAP assay with CANX-EGFP** (Related to Figure 4)

MCF7 cells were transfected with plasmids encoding the CANX-EGFP fusion proteins under control of its cognate 5’UTRs from which the eIF3 binding site was deleted (5’ΔeIF3, **Figure S1B**). After 24 h, live cells were photobleached in the indicated areas of tubular ER or ER whorls, and fluorescence recovery was observed over time. Scale bar: 5 μm.

**Supplementary Movie 3.** **FRAP assay with TMEM33-EGFP** (Related to Figure 4)

MCF7 cells were transfected with plasmids encoding the TMEM33-EGFP fusion proteins under control of its cognate 5’UTRs from which the eIF3 binding site was deleted (5’ΔeIF3, Figure S1B). After 24 h, live cells were photobleached in the indicated areas of tubular ER or ER whorls, and fluorescence recovery was observed over time. Scale bar: 5 μm.

**References**

1.	Duan, H., Zhang, S., Zarai, Y., Öllinger, R., Wu, Y., Sun, L., Hu, C., He, Y., Tian, G., Rad, R., et al. (2023). eIF3 mRNA selectivity profiling reveals eIF3k as a cancer-relevant regulator of ribosome content. The EMBO Journal, e112362. https://doi.org/10.15252/embj.2022112362.

2.	Khatter, H., Myasnikov, A.G., Natchiar, S.K., and Klaholz, B.P. (2015). Structure of the human 80S ribosome. Nature *520*, 640–645. https://doi.org/10.1038/nature14427.
